# Impaired renal gluconeogenesis is a major determinant of acute kidney injury associated mortality

**DOI:** 10.1101/804997

**Authors:** David Legouis, Sven-Erick Ricksten, Anna Faivre, Karim Gariani, Pierre Galichon, Lena Berchtold, Eric Feraille, Kari Koppitch, Alexandre Hertig, Pierre-Yves Martin, Maarten Naesens, Jérôme Pugin, Andrew P McMahon, Pietro E Cippà, Sophie de Seigneux

## Abstract

Acute Kidney Injury (AKI) is strongly associated with adverse outcome and mortality independently of the cause of renal damage^1-3^. The mechanisms determining the deleterious systemic effects of AKI are poorly understood and specific interventions, including optimization of renal replacement therapy, had a marginal effect on AKI-associated mortality in clinical trials^4-8^. The kidney contributes to up to 40% of systemic glucose production by gluconeogenesis during fasting and stress conditions, mainly synthesized from lactate in the proximal tubule^9-12^, rendering this organ a major systemic lactate disposal^13^. Whether kidney gluconeogenesis is impaired during AKI and how this might influence systemic metabolism remains unknown. Here we demonstrate that glucose production and lactate clearance are impaired during human AKI using renal arteriovenous catheterization in patients. Using single cell transcriptomics in mice and RNA sequencing in human biopsies, we show that glycolytic and gluconeogenetic pathways are modified during AKI in the proximal tubule specifically, explaining the metabolic alterations. We further demonstrate that impaired renal gluconeogenesis and lactate clearance during AKI are major determinants of systemic glucose and lactate levels in critically ill patients. Most importantly, altered glucose metabolism in AKI emerged as a major determinant of AKI-associated mortality. Thiamine supplementation restored renal glucose metabolism and substantially reduced AKI-associated mortality in intensive care patients. This study highlights an unappreciated systemic role of renal glucose and lactate metabolism in stress conditions, delineates general mechanisms explaining AKI-associated mortality and introduces a potential therapeutic intervention for a highly prevalent clinical condition with limited therapeutic options.

---

To study the impact of AKI on renal glucose and lactate metabolism, we performed renal vein catheterization in patients undergoing cardiac surgery with cardiopulmonary bypass and experiencing (n=18) or not (n=87) post-operative AKI (**Supplementary Table 1**). We found a switch from net renal lactate uptake to net renal lactate release in patients experiencing AKI (−0.01±0.03 mmol/min to 0.02±0.02 mmol/min, p<0.001). Moreover, AKI patients showed a decrease in the renal net glucose release (0.06±0.07 mmol/min to 0.01±0.04 mmol/min, p=0.016) compared to the control group (**Fig. 1 a,b**). This suggested a simultaneous reduction of gluconeogenesis and activation of glycolysis in the kidney in response to AKI. Glycosuria was unlikely to be involved since all patients had arterial glucose levels below the glycosuria threshold (5.4 ± 0.94 mmol/L). To further characterize this process, we compared kidney allograft biopsies obtained during transplantation shortly (49±16 minutes) after reperfusion in brainstem death donors (n=25), reflecting the early response to ischemic AKI (IRI), to protocol allograft biopsies obtained 12 months after transplantation and considered in recovery state (without ongoing activation of inflammation or fibrosis pathways) (n=17)^14,15^. RNAseq analyses confirmed a reduction of gluconeogenetic and relative increase of glycolytic enzymes mRNA levels in the acute ischemic phase (**Fig 1c**). Living donation implies much shorter ischemia time than deceased donation. When we compared the same pathways in living (n=4) versus brainstem death donor biopsies (n=25), a clear downregulation of gluconeogenesis genes was observed in deceased donors **(Fig. 1d).** Thus, clinical data suggested that gluconeogenesis and glycolysis are modified during AKI, leading to alterations of lactate clearance and glucose production by the kidney.

**Figure 1.**
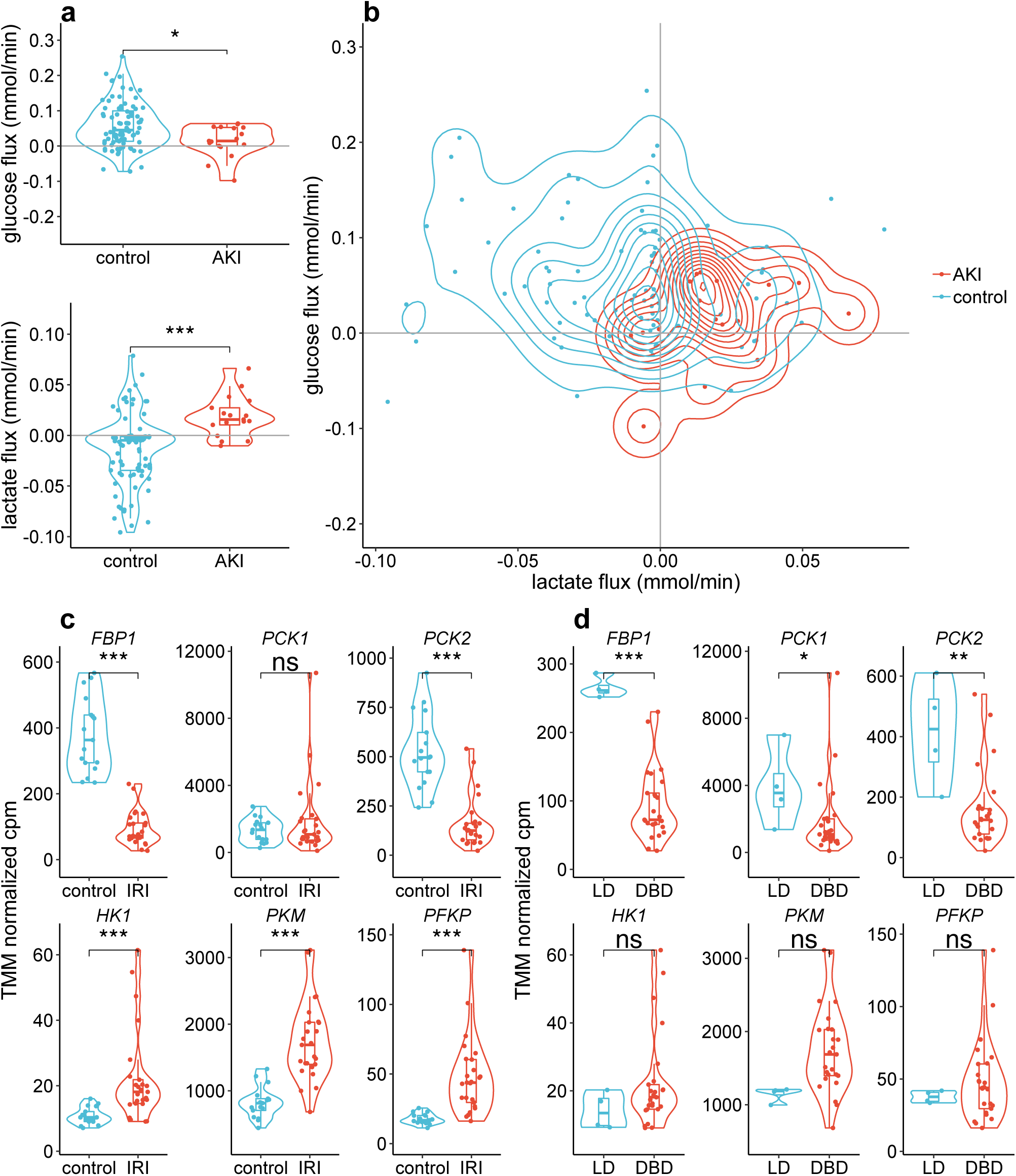
Glycolysis and gluconeogenesis pathway during AKI. (a-b) Glucose and lactate renal flux (mmol/min) in patients undergoing cardiac surgery experiencing or not acute kidney injury (AKI) represented separately in violin plots (smoothed probability density) and boxplots (mean, interquartile, 1st and 95th percentiles) in (a) and in scatter plot with plain lines representing kernel density in (b). The medians were compared using an unpaired Wilcoxon test. (c and d) Expression of gluconeogenesis (FBP1, PCK1, PCK2) and glycolysis (HK1, PKM, PFKP) genes in allograft kidney biopsies obtained at the end of the transplantation from brainstem death donor (IRI group, n=25) and 12 months after transplantation in patients with recovery renal graft status (control group, n=17) (panel d).and comparing the IRI biopsies in kidney recipients patients from living donor (LD, n=4) or donation after brain-stem death (DBD, n=25) (panel d). Differential expression analyses were performed using genewise negative binomial model with quasi likelihood test. *p≤0.05 ; *** p≤0.001

The renal metabolic switch in response to AKI was further investigated in a mouse model of severe ischemia/reperfusion injury (IRI) leading to major tubular damage with denudation of the tubular basement membrane and luminal casts as well as creatinine elevation, as previously reported^16^. Bulk RNAseq analysis revealed a reduction in the expression of enzymes involved in gluconeogenesis (*Fbp1, Pck1*) and a slight increase in the expression of glycolysis enzymes (*Hk1, Pfkp*) during the first hours after IRI, which persisted over the experimental follow-up^16^ (**Fig. 2a,b**). These findings might result from a change in the glucose metabolism in response to ischemia or from a loss of proximal tubule cells by cell death. For a better understanding of this process and avoid extensive tubular necrosis a milder IRI was performed and the tubular response during the injury and the recovery phase was investigated at single cell resolution (**Methods, Extended data Fig. 1,2**). We took advantage of a reporter mouse expressing GFP linked to the histone protein H2B in all cells of the nephron to isolate nephron nuclei after IRI and perform single nucleus transcriptomics (NucSeq) on sorted nuclei (**Fig. 2c, Extended data. Fig. 1,2**). Under steady state, rate limiting enzymes of gluconeogenesis (*Fbp1, Pck1*) were almost exclusively expressed in proximal tubular cells (clusters 0, 1, 2, 7, 13 in **Fig. 2d, Extended data Fig. 1**). In contrast, glycolytic enzymes hexokinase (*Hk1*) and phosphofructokinase (*Pfkp*) were only detectable at low levels in the cells of thick ascending limb of the loop of Henle (cluster 3 in **Fig. 2d**). NucSeq performed on nephron cells harvested at different time points after injury highlighted a widespread response to ischemia in the proximal tubule, characterized by the reduced expression of genes specifically linked to tubular functions, such as membrane transporters (e.g. *Slc34a1, Slc5a12, Slc7a13*), and the up-regulation of genes involved in AKI (e.g. *Havcr1, Gdf15, Krt20*)^16^, cell survival^17^ (e.g. *Clu*) and cell proliferation (e.g. *Cdk6*) (**Fig. 2e, Extended data Fig. 3**). In parallel, we observed a substantial change in the expression profile of genes related to energy metabolism. Focusing on proximal tubule cells, we observed that rate limiting enzymes of gluconeogenesis were strongly reduced during the early phase after IRI (**Fig. 2f,g, Extended data Fig. 3**). The expression level of glycolytic enzymes was slightly increased in proximal tubular cells responding to IRI, but their expression levels remained low. The expression levels of classical transcriptional regulators of gluconeogenesis genes (*PPARA, HNF4*) were downregulated during ischemia reperfusion injury induced AKI in human and mice, and were strongly associated with the expression levels of main gluconeogenesis genes (**Extended data Fig 4**). Twenty-eight days after mild IRI most proximal tubule cells had recovered and displayed a similar transcriptional pattern compared to control mice, including the expression of gluconeogenetic enzymes, but a small subset of cells failed to recover and remained in the non-physiological metabolic state (**Extended data Fig. 2, Fig. 2e**). Consistent with this and the persistent reduction of gluconeogenetic genes in the severe model of IRI (**Fig. 2 a,b**), we confirmed in repeated biopsies from human kidney transplant patients that rate limiting gluconeogenetic enzymes mRNA expression were reduced in CKD and early transition to CKD comparing to biopsies showing evidence for recovery in previously described transcriptional trajectories associated with the transition from AKI to CKD^15^ (**Extended data Fig. 5**). An opposite trend was observed for glycolytic enzymes. Moreover, in human biopsies obtained 3 and 12 months after transplantation lower expression of gluconeogenetic enzymes was associated to a worse one-year renal function, whereas it was not the case in the acute post reperfusion phase (**Extended data Fig. 6 a,b**). Thus, tubular cells go through a fundamental change in energy metabolism in response to AKI. Persistence of these alterations, and particularly impaired gluconeogenesis, is associated to an adverse renal outcome in the long-term.

**Figure 2.**
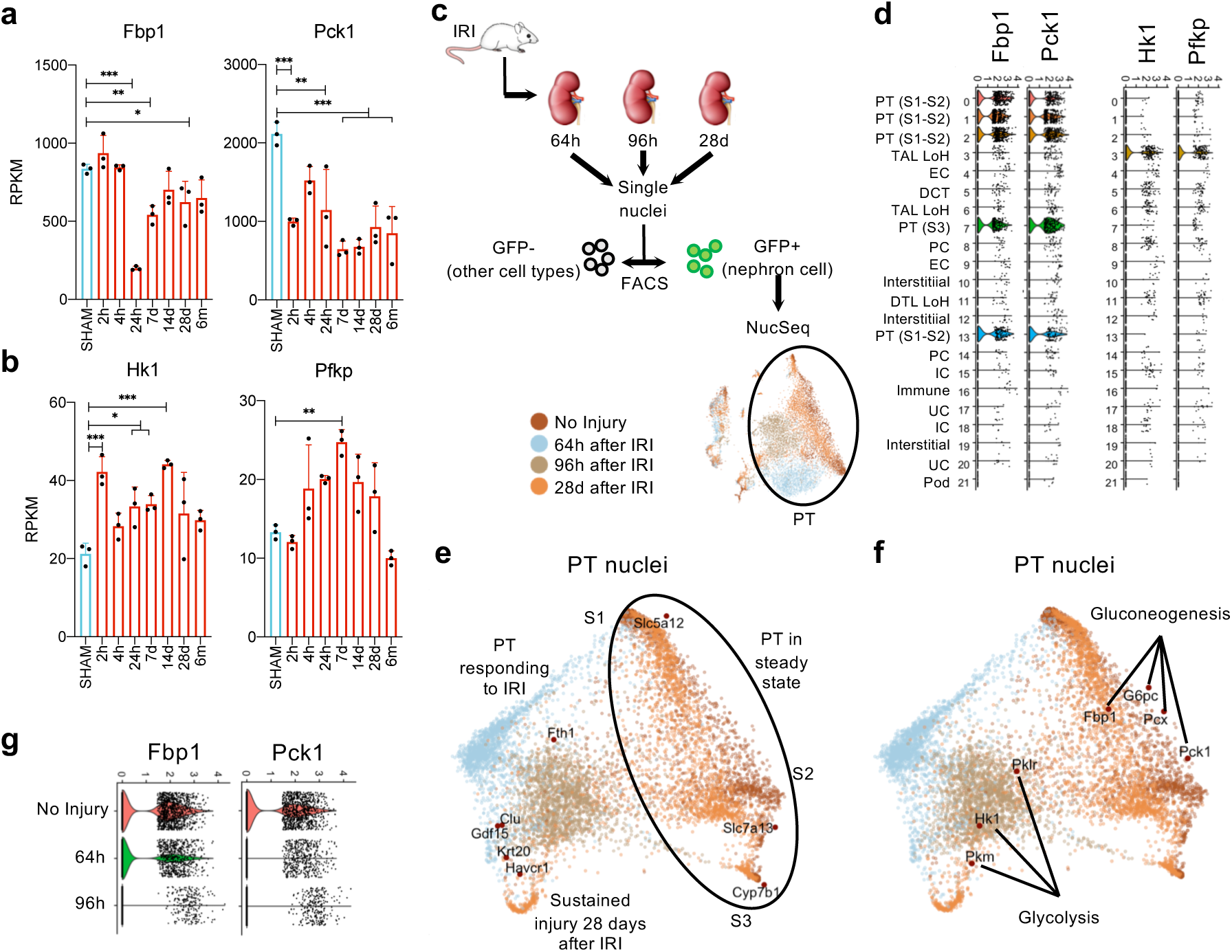
Gluconeogenesis and glycolysis gene expression in the mouse kidney in steady state and in response to IRI. (a-b) RPKM values of representative gluconeogenesis (a) and glycolysis (b) genes in bulk RNAseq in response to 21 minutes IRI (n=3 for each time point). (c) Schematic of experimental strategy; IRI: ischemia/reperfusion injury. (d) Violin plots showing the expression levels of representative gluconeogenesis and glycolysis genes in the cell clusters defined in Suppl. Fig. 2. Cell populations are indicated with initials, as follows: Pod: podocytes, UC: urothelial cells, IC: intercalated cells of the collecting duct, PC: principal cells of the collecting duct, PT: proximal tubule, S1-S2: segment 1 and 2, S3: segment 3, DTL: descending thin limb, LoH: Loop of Henle, EC: endothelial cells, TAL: Thick ascending limb, DCT: distal convolute tubule. (e-f) SWNE plots including PT nuclei isolated at different time points after IRI and highlighting the separation of PT cells in the early phase after IRI from steady state conditions. Representative genes expressed in normal PT cells and in injured cells are embedded in (e). Gluconeogenesis and glycolysis genes are embedded in (f). The color of the dots corresponds to the time point of harvesting after IRI. (g) Violin plot showing the expression level of representative gluconeogenesis genes in PT cells in the early phase after IRI. *p≤0.05 ; **p<0.01 ; *** p≤0.001

In consideration of previous studies linking hypoglycaemia with kidney disease and mortality in hospitalized patients^18-20^, we next explored whether impaired renal gluconeogenesis may result in systemic metabolic alterations in patients with AKI. We analysed simultaneously lactate and glucose levels in 390’803 peripheral arterial blood samples in a cohort of 24’273 intensive care unit (ICU) patients (**Supplementary Table 2**). Lactate levels were higher in patients suffering from AKI. Whereas lactate and glucose levels normally follow the same trend,^21^ AKI patients showed a dissociation at low glucose levels, suggesting a metabolism change **(Fig. 3a).** To further characterize this phenomenon, we categorized each patient according to glucose and lactate levels and we defined following metabolic status : normal glucose/normal lactate (baseline), high glucose/high lactate (stress response), low glucose/normal lactate (isolated low glucose level), high glucose/normal lactate (isolated high glucose level) and low to normal glucose/high lactate (impaired metabolism) **(methods, Extended data Fig. 7**). In multivariable analysis, AKI was the most predictive factor for the low glucose-high lactate combination with a proportionality to AKI severity, as assessed by KDIGO classification (**Fig. 3b**). Propensity score matching was used to abrogate all significant differences between the AKI (n=8’575) and non-AKI (n=17’150) group **(Extended data Fig. 8 and Fig. 9 and Supplementary Table 3**). When comparing the matched groups, the relative number of patients with impaired metabolism pattern was higher in the AKI groups (**Methods, Fig. 3c)**. Moreover, when considering only 495 AKI patients that recovered their renal function during their ICU stay, impaired metabolism pattern was more frequent during AKI than after recovery (**Fig. 3d**). These findings suggest that the kidney is important in determining systemic glucose and lactate levels in ICU patients. In order to exclude a bias related to the association between AKI and general disease severity, we verified this hypothesis in patients with isolated renal ischemia. We measured systemic arterial glucose and lactate in 275 patients admitted to the ICU after kidney transplantation (**Supplementary Table 4**) and compared patients with living (short ischemia time, n=152) or brainstem death (long ischemia time, n=122) donors. In these patients, kidney transplantation surgery was the only reason for ICU admission and IRI can be considered as a primary event and not as a consequence of systemic illness. Patients receiving a kidney from a deceased donor displayed a higher chance of presenting systemic impaired metabolism pattern **(Fig 3e)**, in line with the mRNA regulations in biopsies from kidney allograft patients (**Fig. 1 c,d**). In addition, we observed a correlation between impaired metabolism status and 15 days creatinine levels. (**Extended data Fig. 10**). Thus, kidney injury is an important and independent determinant of systemic glucose and lactate levels. Next, we investigated the impact of systemic alterations of lactate clearance and glucose production on mortality in ICU patients. As expected, AKI was associated with increased mortality, but this effect was restricted to patients with impaired metabolism pattern after propensity score matching (**Fig. 3f and Extended data Fig. 11)**. These findings indicate that alterations of systemic metabolism have a stronger impact on patient mortality than AKI, but that AKI is a major determinant of such systemic metabolic alterations. Thus, the well-known association between AKI and mortality might be secondary to the systemic impact of the alteration in lactate and glucose metabolism in the proximal tubule.

**Figure 3.**
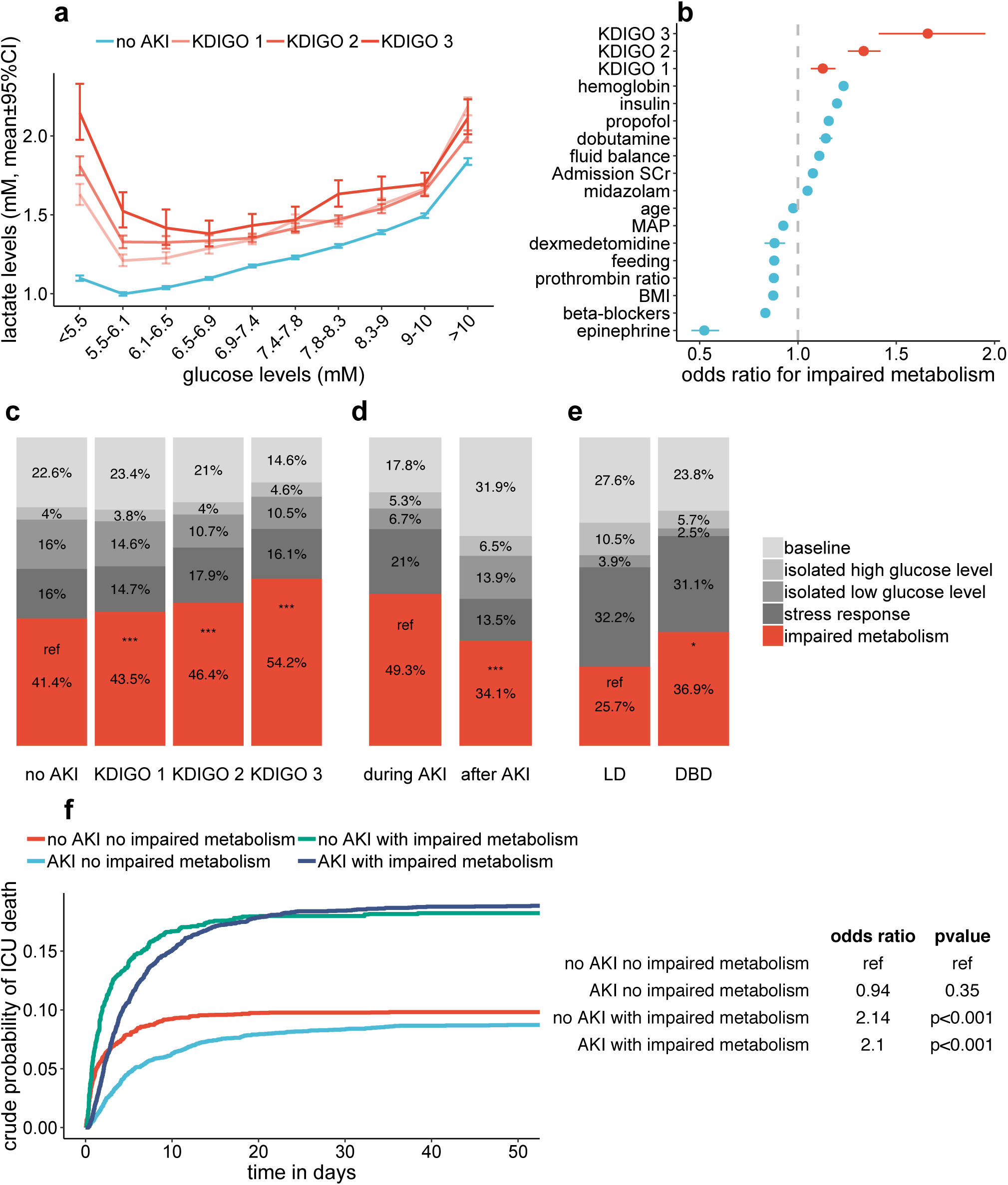
Impact of AKI on systemic metabolism and ICU mortality. (a) Scatter plot showing the relation between all the lactate levels recorded in the ICU datasets and glucose levels, stratified according to KDIGO status (n=390’803). For each glucose level range, mean ± 95% CI was plotted. (b) Odds ratio with 95% confidence interval for having impaired metabolism status (low glucose, high lactate) obtained from multivariate logistic linear mixed model. (c,d) Stacked barplots showing metabolism pattern: (c) in ICU patients matched using a propensity score for AKI (n= 8575 for AKI and n=17150 for non AKI patients), (d) in ICU patients experiencing AKI (n=495), during the episode and after its resolution and (e) in allograft kidney recipients from living (LD, n=152) or brain death donor (DBD, n=122). Comparisons were performed using a conditional logistic model stratified on patients or matched patients. (f) Cumulative incidence curves for ICU mortality in the same cohort of ICU patients matched using a propensity score for AKI in 4 subgroups, with or without AKI and with or without impaired metabolism assessed using a conditional logistic regression.

If this hypothesis is correct, restoring renal gluconeogenesis and limiting glycolysis is expected to modify the dismal prognosis of critically ill patients with AKI. Thiamine plays an essential role in oxidative metabolism^22^ and thiamine deficiency was associated with altered glycolysis and gluconeogenesis in rats and mice.^23^ Thiamine deficiency occurs frequently in critical care patients.^24^ Consistently with previous studies, ^23,25^ thiamine decreased glycolysis rate in a dose-dependent manner in renal tubular cells in vitro (**Fig. 4a**). In a fresh suspension of isolated mouse proximal tubular cells, thiamine enhanced glucose production (**Fig. 4b**). Therefore, we reasoned that thiamine supplementation might partially compensate the systemic alterations associated to AKI. In our ICU, thiamine is administered in patients with suspected vitamin deficiency (e.g. in the context of alcoholism or malnutrition) and more extensively in septic patients^23,26^, thereby offering a clinical model to study our hypothesis. In a retrospective analysis, we matched 347 ICU AKI patients receiving high dose thiamine (cumulative dose > 500 mg) to 694 patients who did not, using a propensity score (**Extended data Fig. 12, Supplementary Table 5**). All the considered variables were balanced between groups (**Extended data Fig. 13**). In patients with AKI, thiamine administration was associated with a rapid decrease in the impaired metabolism pattern **(Fig. 4c).** Most importantly, and consistently with our hypothesis, their mortality was significantly reduced (OR for mortality 0.59 (0.14-0.86). **(Fig. 4d)**. The same matching strategy was used to compare 263 ICU patients without AKI but receiving thiamine supplementation and 526 patients who did not. Neither metabolism pattern **(Fig. 4e)** nor mortality **(Fig. 4f)** were significantly affected by thiamine treatment in patients without AKI, although a trend was observed for mortality.

**Figure 4.**
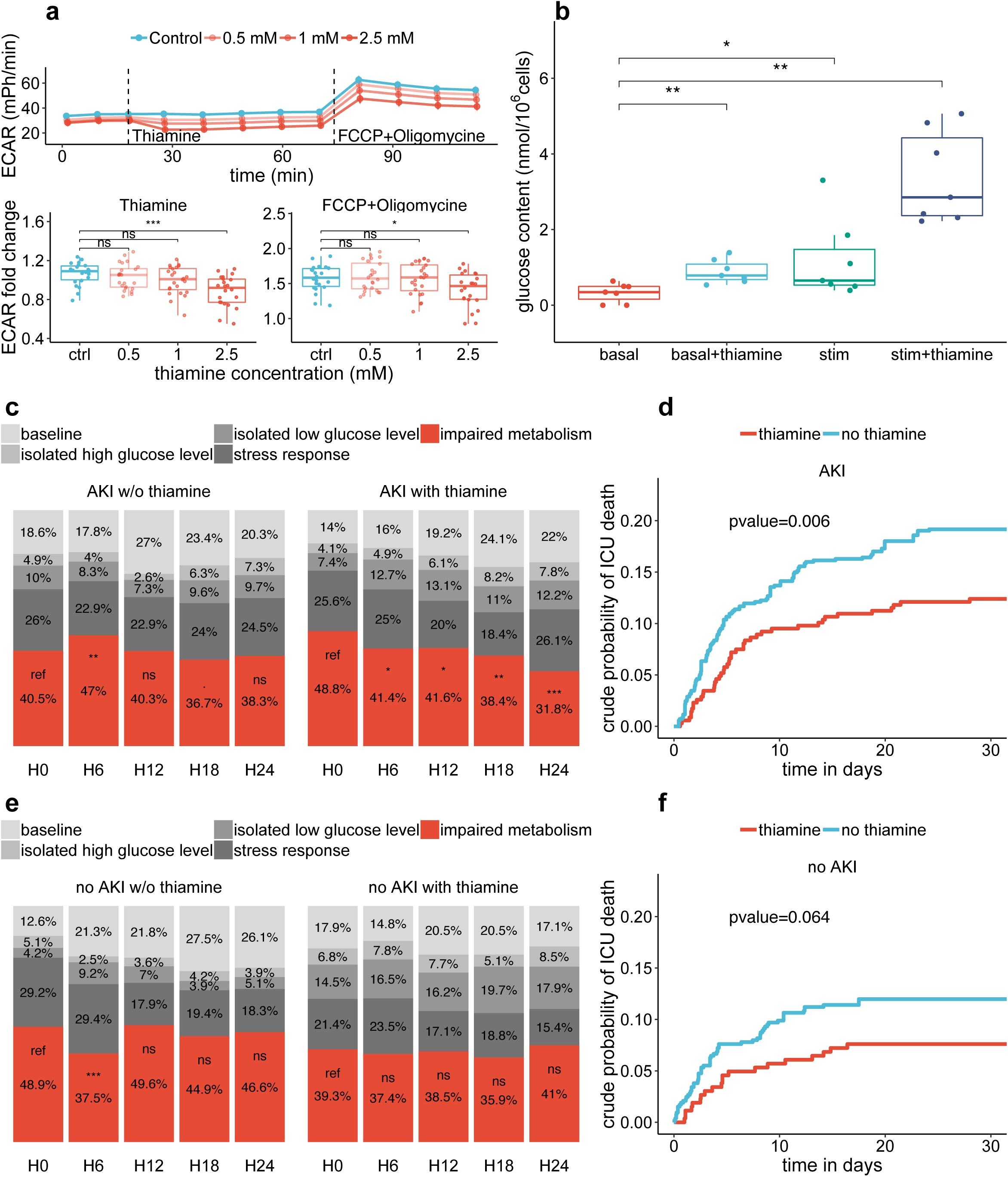
Effect of thiamine supplementation on glucose metabolism and mortality. (a) Extracellular acidification rate (ECAR) of HK2 cells measured in a Seahorse XF96 analyzer using oligomycin and FCCP mixture as stress test and 3 different concentrations of thiamine added to the medium after 18 minutes of stabilization. Data were showed as mean ± 95% confidence interval (top panel) and as fold change in baseline ECAR (bottom panel). (b) glucose content measured in supernatant of a fresh suspension of primary isolated proximal tubular cells and incubated for 120 minutes with basa medium (KRBH), KRBH and thiamine (basal+thiamine), stimulation medium (KRBH with 10 mM pyruvate, 10 mM lactate and 10 mM glutamine) or stimulation medium with 2.5 mM thiamine (stim+thiamine). Data are normalised on number of cells. (c) Stacked barplots showing metabolism pattern over time in ICU patients experiencing AKI and receiving (n=347) or not (n=694) thiamine supplementation according to a propensity score matching. (d) Cumulative incidence curves for ICU mortality in the same AKI patients matched according to their thiamine supplementation status. (e) Stacked barplots showing metabolism pattern over time in ICU patients free of AKI and receiving (n=263) or not (n=526) thiamine supplementation according to a propensity score matching. (d) Cumulative incidence curves for ICU mortality in the same non-AKI patients matched according to their thiamine supplementation status. Comparisons were performed with multivariate conditional logistic regression stratified on matched patients and generalised linear mixed model. *p≤0.05 ; **p≤0.01 ; *** p≤0.001

Altogether, our data highlight the pivotal role of the kidney as a metabolic organ and reveal an unappreciated effect of renal gluconeogenesis in regulating systemic glucose and lactate levels in conditions of acute stress. We demonstrate that impaired renal gluconeogenesis during AKI alters systemic metabolic homeostasis, playing a major role in AKI-associated mortality, and is accessible to a therapeutic intervention.

## Methods

### Renal catheterization during human AKI

Renal vein catheterization was performed in postcardiac surgery patients as previously described^27-29^. After placement of the catheter in the left renal vein, renal blood flow was measured by the retrograde thermodilution technique. For measurement of the glomerular filtration rate, an intravenous priming dose of the filtration marker, ^51^Cr-EDTA, was given, followed by an infusion at a constant rate individualized to the body weight and serum creatinine. Arterial and renal vein blood samples were obtained twice with a 30-minutes interval. In the non-AKI group, measurements were performed 4-6 hours after end of surgery. In the AKI group, measurements were performed 2-6 days after surgery. All patients were sedated and mechanically ventilated. The study was performed according to Helsinki’s principles and was approved by the human ethics committee of the University of Gothenburg. AKI was defined by KDIGO criteria^30^. Patient’s clinical characteristics are described in **supplementary table 1**.

Renal Arterial Blood Flow,

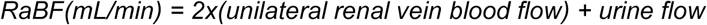

Renal Venous Blood Flow,

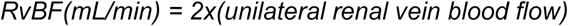

Glucose and lactate renal net flux,

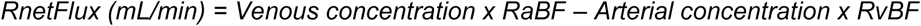

### Ischemia reperfusion injury in mice

Mouse handling, husbandry and all surgical procedures were performed according to guidelines issued by the Institutional Animal Care and Use Committees (IACUC) at University of Southern California. The experimental model was previously characterized.^16^ Briefly, 10-to 12-week-old, 25-28g, male mice, were anesthetized with an intraperitoneal injection of a ketamine/xylazine (105 mg ketamine/kg; 10 mg xylazine/kg). Body temperature was maintained at 36.5-37°C throughout the procedure. The kidneys were exposed by a midline abdominal incision and both renal pedicles were clamped using non-traumatic microaneurysm clips (Roboz Surgical Instrument Co.). Ischemia time was determined as indicated below. Restoration of blood flow was monitored by the return of normal color after removal of the clamps. All the mice received intraperitoneal (i.p) 1 ml of normal saline at the end of the procedure. Sham-operated mice underwent the same procedure except for clamping of the pedicles. Ethynyldeoxyuridine (EDU) (0.05mg/g, Sigma-900584) was injected 48h after surgery to be able to identify cells proliferating after injury. Serum creatinine levels were measured at the UT Southwestern Medical Center O’Brien Center for Kidney Disease Research by capillary electrophoresis (PA800 Plus Pharmaceutical Analysis System, Beckman Coulter). BUN was assayed on the Stanbio Excel analyzer using BUN Procedure 0580 for the quantitative colorimetric determination of urea nitrogen in serum at George M. O’Brien Kidney Center at Yale University.

Ischemia time was specifically determined in two sets of experiments: First, ischemia time was optimized to obtain the most severe renal injury allowing long-term mouse survival, as previously reported^15^. The best ischemia time for this purpose was determined to be 21 min. In a second set of experiments, the ischemia time was reduced to achieve substantial tubular injury in the early phase, but almost complete recovery during follow-up (**suppl. Fig. 2**): the level of injury was determined 48h after surgery, the level of repair 28 days after surgery by measuring serum creatinine and BUN, by histology and by qPCR. The best ischemia time for this purpose was determined to be 15 min.

### Histology and immunofluorescence

For conventional staining kidneys were perfused with ice-cold PBS and embedded in paraffin after overnight fixation in 4% paraformaldehyde (PFA) at 4°C. Sections were cut at 2 µm and stained with hematoxylin and eosin. For immunofluorescence PFA fixed tissues were equilibrated in 30% sucrose/PBS overnight then embedded in OCT in dry ice ethanol bath. 8-10 µm frozen sections were washed in PBT (PBS + 1% Triton-X), blocked in in 5% Normal Donkey Serum in PBT, and incubated overnight at 4°C with primary antibodies and detected with species-specific secondary antibodies coupled to Alexa Fluor 488, 555, 594 and 647 (Life Technologies) for 1 hour at room temperature. Following antibodies were used in this study to recognize Havcr1 (R&D Systems AF1817, 1:500), LTL (Vector Laboratories FL-1321, 1:250), UMOD (Thermo Fisher J65429, 1:250), AQP2 (Santa Cruz sc-515770 1:500). All sections were stained with Hoechst 33342 (Life Technologies) prior to mounting with Immu-Mount (Fisher). All images were acquired on Zeiss Axio Scan Z1 slide scanner and Zeiss LSM780. EDU staining on sections was performed with Click-iT™ Plus EdU staining according to the manufacturer’s protocol (Thermo C10640).

### Nephron nuclei isolation and sorting

To investigate single cell transcriptomics on renal tubule cells, we generated a reporter mouse expressing GFP linked to the histone protein H2B in all nephron cells by combining 3 transgenes (Six2TGC;Rosa26rtTA;pTREH2-GFP; shortly Neph-GFP). Neph-GFP mice express a Cre recombinase under the control of Six2 in nephron progenitor cells during development (Six2-TGC),^31^ which activates the tetracycline-controlled transcriptional activator (rtTA) under the control of the constitutively active Rosa26 promoter (Rosa26rtTA). Six2 is not active in the adult kidney, but all cells derived from Six2+ progenitors continue to synthetize the rtTA, which activates the tetracycline response element (TRE) leading to GFP-H2B expression in the presence of doxycycline, independently of gene transcription regulation. Neph-GFP mice were kept on doxycycline food (purchased from Envigo TD.08541) starting a few days after birth. The validation of the mouse model is presented in **suppl. Fig. 1**. For a single-cell transcriptional profile, at time of harvesting animals were perfused with ice cold HPBS till kidneys cleared of blood. Kidney halves were immediately placed in cold HPBS with RNasin Plus RNase Inhibitor (0.4U/µl, purchased from Promega N2615) and cut into small pieces (>25). These pieces were immediately transferred into a 2 ml dounce (D8938 SIGMA) loaded with 1ml of NEZ Lysis Buffer (+ Rnase inhibitor) on ice. Samples were then dounced on ice with 5 strokes of the looser pestle (A) every 2 minutes for 8 mins (25 strokes total). Samples were then dounced 25 times slowly with the tighter pestle (B) on ice. The dounced sample was then filtered through a 40-µm Falcon Nylon Cell Strainer. The filter was then washed with 8 ml of 1% BSA PBS (diluted from SRE0036 SIGMA in CaCl/MgCl free HPBS + **Rn**ase Inh) and nuclei suspension spun in a precooled (4°C) centrifuge at 650g for 8 min. Supernatant was removed and pellet was re-suspended in 1%BSA PBS (+Rnase Inh) and moved to a low bind Eppendorf tube. Nuclei quality and number were assessed with Tryptan Blue staining and a hemocytometer. An appropriate number of nuclei was then pipetted out and spun down again (700g/8min). Supernatant was removed and pellet was resuspended in 2%BSA (+RNase inhibitor) by excessive pipetting to avoid clumping and kept on ice until FACS sorting. Nuclei were sorted on a BD Aria 1 into same 2% BSA solution. The sorting strategy is presented in **Suppl. Fig. 1b**: nuclei were first sorted as DAPI positive and then for GFP expression.

### NucSeq and data analysis

In total 8 samples were processed and included in following analyses, including GFP+ nuclei no surgery, GFP-nuclei no surgery, GFP+ nuclei harvested 64h after IRI and sham surgery, GFP+ nuclei harvested 96h after IRI and sham surgery, GFP+ nuclei harvested 28 days after IRI and sham surgery. After sorting nuclei were counted and immediately processed according to 10X Chromium protocol. Briefly, an appropriate volume of each cell suspension that gave 7,000 - 9,000 cells was combined with freshly prepared 10X Chromium reagent mix and the samples were loaded into separate lanes of a microfluidic partitioning device, per the 10X Genomics Chromium v2 Single Cell 3’ reagent kit instructions. Cell capture, lysis and mRNA reverse transcription occurred in-droplets.cDNA recovered from the emulsion was cleaned-up, amplified by PCR, checked for size and yield on a 4200 Tape station (Agilent), processed into barcoded Illumina ready sequencing libraries and again assayed for size and yield. The Translational Genomics Center at Children’s Hospital Los Angeles Center for Personalized Medicine carried out paired-end sequencing on the HiSeq 4000 platform (Illumina) using the HiSeq 3000/4000 SBS PE clustering kit (PE-410-001) and 150 cycle flow cell (FC-410-1002) and processed raw data into fastq files. Alignment of sequencing reads to the mouse pre-mRNA reference genome, as well as generation of BAM files and filtered gene-barcode matrices was accomplished by running Cell Ranger Single-Cell Software Suite 2.0 (10X Genomics) using the STAR aligner on the USC High Performance Cluster. The Cell Ranger *cellranger count* function output filtered gene-cell expression matrices removing cell barcodes not represented in cells. Principle component analysis and identification of variably expressed genes were carried out using the Seurat package v 2.3.4 in R Studio. Briefly, for the primary analysis, the matrices from the individual samples were first merged as required into a metadata annotated r-object using the *Read10X, CreateSeuratObject AddMetaData* and *MergeSeurat* functions. The raw dataset was then filtered with range filters for genes/cell (400-3,000). The resulting primary analysis dataset contained 6,086 cells for the dataset including GFP+ and GFP-nuclei from mice without injury and 11’274 cells for the dataset including GFP+ nuclei from mice harvested at different time points after IRI. The *NormalizeData* function scaled and log transformed that dataset. Following the *FindVariableGenes*, the principle component analysis first on the variable genes with *RunPCA* was performed. The useful principle components were used in graph-based clustering using *FindClusters* at resolution 1.2. *FindAllMarkers* generated the list of genes differentially expressed in each cluster compared to all other cells based on the Wilcoxon rank-sum test and limiting the analysis with a cutoff for minimum log FC difference (0.25) and minimum cells with expression (0.25). The secondary Seurat analyses on PT cells employed the *SubsetData* function to create new r-objects from cohorts of primary analysis. For data visualization we used *RunTSNE* followed by *TSNEPlot, FeaturePlot, DotPlot* and *VlnPlot* function in Seurat. To better characterize the PT subset, we used the SWNE (Similarity Weighted Nonnegative Embedding) package v 0.2.21 in R Studio^32^. Starting from Seurat objects we applied performed SWNE embedding with following parameters: *k* = 12, *alpha.exp* = 1.9, *snn.exp* = 0.1, *n_pull* = 3. Gene of interested were embedded into the SWNE plots to enhance biological interpretation.

### Renal allograft RNA sequencing

Genome wide gene expression profile using RNA sequencing was performed in kidney allograft recipients as previously described^15^. Briefly 42 patients were enrolled at the University Hospitals of Leuven. For each of them, protocol biopsy was performed at four different timepoints: before implantation (kidney flushed and stored in ice), after reperfusion (at the end of the surgical procedure) and 3 months and 12 months after transplantation. Raw genes counts were normalized by the trimmed mean of M-values method (TMM) using edgeR package and genes with null expression in one sample were excluded. Counts were expressed in counts per million (cpm). Relation between continuous clinical outcomes and mRNA expression were fitted using a robust linear model. Recovery biopsies were identified as previously described using a transcription analysis^15^. Biopsies performed at the time of reperfusion and issued from donors after cardiac death (n=13) were excluded from the analysis. Comparisons of genes expression were performed using genewise negative binomial linear model with quasi likelihood test.

### Clinical analysis of intensive care unit patients

In a retrospective study, we extracted all available Clinical, Therapeutic and Laboratory data from all patients admitted to the ICU of the University hospital of Geneva from 2007 to 2018 using SQL request exported to an anonymized csv file.

#### Metabolism status

Each glucose and lactate levels pair available in the long dataset were classified in the 5 following groups. baseline (lactate levels below median and with glucose levels between the 25^th^ and the 50^th^ percentile) ; impaired metabolism (lactate levels above the median with glucose level below the 75^th^ percentile) ; isolated low glucose level (lactate levels below median with glucose levels above the 75^th^ percentile) ; isolated high glucose level (lactate levels below median with glucose levels below the 25^th^ percentile) and stress response (lactate levels above median and glucose levels above the 75^th^ percentile). These groups defined the metabolism status associated to each glucose and lactate levels pair.

#### Metabolism pattern

Metabolism pattern was defined as the most represented metabolism status during the whole ICU stay for each non-AKI and kidney allograft patients. For each AKI patient, the most represented metabolism status was chosen according to the status during or after AKI.

*Acute Kidney Injury (AKI)* was defined by an 1.5-fold increase in first serum creatinine levels measured in the ICU or a urine output less than 0.5 mL/kg/h during at least 6 consecutives hours; stage 2 was defined as increase in creatinine above 2-fold from first serum creatinine levels or urine output less than 0.5 mL/kg/h during at least 12 consecutive hours; stage 3 was defined as above 3-fold from first serum creatinine levels, or any serum creatinine >4 mg/dL or urine output less than 0.5 mL/kg/h during at least 24 consecutive hours.

#### During and after AKI status

As serum creatinine levels was measured several times, we defined, for AKI patients two AKI time status: during AKI as long as serum creatinine levels was above 80% of the maximal recorded value during the ICU stay for the considered patient and after AKI if serum creatinine levels drop below 120% of the serum creatinine levels at ICU admission.

*Missing values* were managed depending on how frequently they were missing. Variables with missing values above 5% were imputed by bagged tree whereas those with missing values below 5% were analysed in complete cases. No variables had missing values rate above 20%.

#### Data management

Long datasest contained all the glucose-lactate levels pairs measured, one per row and all the associated variables available, recorded at the closest time. Short datasets were aggregated by patients. Numerical associated variables were summarised by median and therapeutic variables by Boolean values. For AKI patients, two datasets were built with the same procedure, depending or not of AKI time status. Similarly, a short and a long thiamine dataset were built but including only patient admitted between 2016 January 1^st^ and 2018 December 31^st^.

#### Propensity score calculations

Two propensity scores for AKI and thiamine supplementation were estimated with a non-parsimonious logistic regression using AKI and thiamine short datasets respectively. The variables included were SAPS score, sex, age, body mass index, haemoglobin, insulin, feeding prothrombin ratio, bilirubin, eGFR at ICU admission, dobutamine, epinephrine, norepinephrine, terlipressin, midazolam, propofol, metformin and betablockers for both AKI and thiamine datasets. For thiamine, we fitted propensity score on split datasets according to the AKI status.

#### Propensity score matching

The two propensity scores estimated were used to match each patient developing AKI or receiving thiamine supplementation to two controls with similar logits of the propensity score. We used matching strategy based on nearest neighbour matching with replacement (controls can be used more than once) using callipers of width equal to 0.1 SD. An absolute standardized difference less than 10% was considered to support the assumption of balance between groups. The standardized differences for all variables used are showed in extended data.

Lactate versus glucose levels plot was performed using the AKI long dataset. For the lactate-glucose levels pairs from AKI patients, glucose and lactate levels should have measured during AKI only and not after. 95% confidence interval were estimated using bootstrapping.

Association between available variables and impaired metabolism status was studied using a mixed logistic regression and the AKI long dataset. The numerical variables were first scales, centred and transformed by Yeo-Johnson method to make them more normal like. It allows easiest comparisons of odds ratio with logical variables as mean was equal to 0 and 1 correspond to the 85^th^ percentile of the considered variable. Random effect was defined by patient in the form 1|patient. The 95% confidence intervals for odds ratios were estimated using modified Wald method.

Comparisons of metabolism pattern was performed on the AKI matched short dataset for the KDIGO status and on the AKI short dataset for the during and after comparisons. For the kidney allograft recipients’ analyses, we excluded patient with combined transplantation to avoid confusion factors. Deceased donors were only brain death donor. For the time course analyses after thiamine supplementation, we used matched patients from thiamine matched dataset. For each matched patients we extracted the exact delay between ICU admission and thiamine supplementation. This delay was used as start time for time course analyses for both thiamine recipients and control patients. We then recalculated for each patient the metabolism pattern at 5 different time points (H0, H6, H12, H18 and H24) and analyses the occurrence of impaired metabolism pattern using logistic mixed model with random effect for matched patients.

Mortality analyses was performed on matched AKI and thiamine datasets and differences between groups were assessed using conditional logistic regression stratified on matched patients.

Data mining and statistical analysis were performed using R software. The study was approved by the local ethical committee for human studies of Geneva, Switzerland (CCER 201-00320, Commission Cantonale d’Ethique de la Recherche) and performed according to the Declaration of Helsinki principles.

### Cell energy phenotyping under thiamine treatment

An assay using the Seahorse XF-96 Extracellular Flux Analyzer (Agilent) was performed to measure the acidification rate. Briefly, single cell solution of HK2 was plated in XF96 Cell Culture Microplates (Seahorse Agilent) at a cellular density of 40,000 cells/well. On the time of the experiment, cell media was replaced by Seahorse assay media (DMEM medium supplemented with glucose 10 mM, pyruvate 1 mM and L-Glutamine 2 mM) and cells were incubated at 37°C in a carbon dioxide-free incubator for 1 hour before assessing basal glycolysis rate. Thirty minutes before the experiment, the sensor was placed into the XF-96 instrument and calibration was initiated with XF calibrant medium (Agilent). After calibration, the basal extracellular acidification rate was recorded for 20 minutes. Thiamine was injected using Seahorse ports and the acidification rate was recorded during 55 minutes. Then, FCCP (2 µM) and Oligomycin (1 µM) were co-administrated to assess metabolic potential and the ECAR was recorded for 45 minutes. We used 24 replicates in each condition. Three experiments were performed. The person who analyzed the data was blinded to the experimental treatment, but not the experimenter. pH of all media including thiamine were adjusted to 7.4.

### Production of glucose in primary mouse proximal tubular cells

Primary mice proximal tubular cells were obtained as previously described^33^. Briefly mice were scarified and the kidneys harvested. They were then mechanically dissociated using the GentleMACS cell dissociator (Miltenyl Biotec, California, USA). The proximal tubular cells were isolated with anti Prominin1 microbead conjugated antibodies and autoMACS (Miltenyl Biotec, California, USA). Cells were further incubated for 120 minutes at 37°C in an incubator with 5% CO2 in four different media: KRBH only, KRBH with thiamine 2,5 mM, KRBH with, 10 mM pyruvate, 10 mM lactate and 10 mM glutamine and KRBH with 10 mM pyruvate, 10 mM lactate and 10 mM glutamine and 2.5 mM thiamine. Supernatant was taken for glucose quantification with glucose assay kit (Biovision) and later normalized by number of cells.

### Statistical analysis

Baseline characteristics were expressed as mean (standard deviation) and median (25-75^th^ percentiles) or absolute and relative (%) frequency if categorical. They were compared by Student’s t, Mann Whitney or Chi-square tests depending on their class and their distribution.

Data from renal venous catheterization were compared using Mann Whitney test.

Normality of the residuals and homoscedasticity were assessed using a qq-plot and scatter plot respectively. Variables with a univariate *P*-value <0.5 were first considered in the multivariable analysis, as well as clinically pertinent interaction between variables, and the final regression model was performed using modified backward selection. Absence of multicollinearity was checked against a correlation matrix.

A p-value of less than 0.05 was considered significant and all p-values were two-tailed and adjusted for multiples comparisons using Benjamini-Hochberg correction if necessary.

## Data availability

RNAseq data for human kidney biopsies are respectively available at GEO GSE126805. Mouse RNAseq data are available on GEO, PMID: 24569379 [https://www.ncbi.nlm.nih.gov/geo/query/acc.cgi?acc=GSE52004] and as supplementary table at https://doi.org/10.1172/jci.insight.94716. NucSeq data will be made available upon publication. Clinical data are available from the corresponding authors upon reasonable request. Raw data are included in this published article (and its supplementary information files).

## Code availability

Custom code will be made available on gitlab.unige.ch upon publication.

## Acknowledgements

This study is funded by the Swiss National Science Foundation (grant PP00P3 157454 to Sophie de Seigneux and grant 167773 to Pietro Cippà), by a STARTER grant (RS03-25) to Sophie de Seigneux and David Legouis from the HUG private foundation (the foundation of the Geneva University Hospitals and the University of Geneva’s Faculty of Medicine) and by The Eli and Edythe Broad Foundation (Pietro Cippà and Andrew McMahon). We thank Greg Alvarado, Andy Ransick and Albert Kim for technical support.

## Extended data figure legends

**Extended data figure 1**

Validation of the Neph-GFP mouse. (a) Schematic representation of the separation in nephron and non-nephron cells in Neph-GFP mice. (b) Representative example of flowcytometry and sorting of nuclei isolated from the kidney of a Neph-GFP mouse. (c-d) TSNE analysis of merged GFP^+^ and GFP^-^cells from one representative Neph-GFP kidney. Cell clusters are labelled with numbers and cell populations with initials in (c) (s. Fig. 2d), the separation in GFP^+^ and GFP^-^cells is shown in (d). Please note that a subset of nephron cells was detected in the GFP^-^fraction, probably as a result of the delay in the replacement of histone proteins. The presence of cells with a transcriptional profile overlapping with principal and intercalated cells of the collecting duct in the GFP^+^ fraction was consistent with previous observations. (e) Dotplots of representative marker genes for each cell cluster. Cell clusters are labelled with numbers, cell populations with initials, as follows: Pod: podocytes, UC: urothelial cells, IC: intercalated cells of the collecting duct, PC: principal cells of the collecting duct, PT: proximal tubule, S1-S2: segment 1 and 2, S3: segment 3, DTL: descending thin limb, LoH: Loop of Henle, EC: endothelial cells, TAL: Thick ascending limb, DCT: distal convolute tubule. (f-h) Immunofluorescence from a representative example of a Neph-GFP mouse kidney showing GFP^+^ nuclei in Ltl^+^ PT cells and Umod^+^ TAL cells, but GFP^-^nuclei in Aqp2^+^ collecting duct cells.

**Extended data figure 2**

A mouse model of moderate IRI to study tubular repair. (a) Serum creatinine values measured 48h after IRI, with different ischemia times, as indicated. N=3-6 mice. (b) BUN values measure 48h and 28 days after IRI (15 minutes ischemia). N=5-12 mice. (c) qPCR on renal tissue isolated at different time points after IRI with different ischemia times, as indicated. N=5-8 mice, **P<0.01. (d-g) Conventional H&E staining (d, f) and immunofluorescence (e, g) on kidney sections obtained 2 and 28 days after 15 min IRI showing the presence of damaged tubular cell at the cortico-medullary junction (S3 region of the PT). 28 days after IRI most tubular cells recovered, but still retained EDU (injected 48h after IRI), but some flattened cells and damaged tubules persisted (f).

**Extended data figure 3**

NucSeq analysis after IRI. (a) TSNE analysis on merged datasets including GFP+ nuclei at different time points after IRI as indicated. The line delineates proximal tubule (PT) cells. S1 and S3 indicate the corresponding segments of the PT. (b) Feature plots of representative genes, typically expressed by proximal tubule cells (*Slc34a1*), S1 cells (*Slc5a12*), S3 cells (*Cyp7b1*) and injured proximal tubule cells (*Fth1*). (c) SWNE plot including PT nuclei isolated at different time points after IRI as presented in Fig. 2. The colors indicate the expression level of representative genes as indicated.

**Extended data figure 4**

mRNA expression of gluconeogenesis regulators, *HNFA4* and *PPARA*. (a) TMM normalised cpm of HNFA4 and PPARA genes in allograft kidney biopsies obtained at the end of the transplantation from brain-stem death donor (IRI group, n=25) and 3 or 12 months after transplantation in patients with recovery renal graft status (control group, n=23). (b) relation between *HNFA4* and *PPARA* genes with gluconeogenesis genes *FBP1, PCK1 PCK2* and *PC.* (c) SWNE plots including PT nuclei isolated at different time points after IRI and highlighting the separation of PT cells in the early phase after IRI from steady state conditions. Representative regulators genes expressed in normal PT cells are embedded The color of the dots corresponds to the time point of harvesting after IRI. Violin plot and boxplot (with mean, IQR, 1^st^ and 95^th^ percentiles) are shown. Differential expression analyses were performed using genewise negative binomial model with quasi-likelihood test. Relation between genes expression was modelised using linear model. *p≤0.05; *** p≤0.001

**Extended data figure 5**

FBP1, PCK1, PCK2, HK1, PFKP and PKM genes expression data from RNAseq performed 3 and 12 months after transplantation and classified as recovery (red, n=23), early transition to CKD (green, n=22) and CKD (blue, n=27) using machine learning computational approach^15^.

**Extended data figure 6**

Relation between glycolytic and gluconeogenic gene expression and one-year renal function in kidney allograft recipients (a) Relation between FPB1 (left panel), *PKM* (right panel) genes expression at 3 months after transplantation and one-year GFR estimated by the CKD-EPI equation. The regression was fitted using a robust linear model. (b) Relation between one-year eGFR and *FBP1, PC, PKM* and *PFKP* gene expression at 3 different timepoints (during the transplantation, 3 and 12 months after) in kidney allograft recipients. Each dot represents the coefficient (±95% CI) of the robust linear model fitting the relation between gene expression levels and one-year eGFR. Positive value indicates a positive association.

**Extended data figure 7**

Scatter plot showing all the glucose and lactate values recorded in the ICU datasets (n=661’557). Five metabolism status were defined : baseline (lactate levels below median and with glucose levels between the 25^th^ and the 50^th^ percentile) ; impaired metabolism (lactate levels above the median with glucose level below the 75^th^ percentile) ; isolated low glucose level (lactate levels below median with glucose levels above the 75^th^ percentile) ; isolated high glucose level (lactate levels below median with glucose levels below the 25^th^ percentile) and stress response (lactate levels above median and glucose levels above the 75^th^ percentile).

**Extended data figure 8**

Relative standardized differences between AKI and non AKI patients, before and after propensity score matching for each variable included in the propensity score. A relative standardised difference less than 10% was considered to support the assumption of balance between groups.

**Extended data figure 9**

Flowchart of the propensity score matching strategy

**Extended data figure 10**

Relation between proportion of metabolism status classified as impaired metabolism, for each allograft kidney recipients during the ICU stay and the post-operative day 15 serum creatinine levels, assessed by robust linear model

**Extended data figure 11**

Cumulative incidence curves for ICU mortality in the cohort of ICU patients matched using a propensity score for AKI comparing 4 groups (with or without impaired metabolism pattern and with or without AKI) stratified for KDIGO.

**Extended data figure 12**

Flowchart of the propensity score matching strategy

**Extended data figure 13**

Relative standardized differences between patients receiving or not thiamine supplementation, with or without AKI, before and after propensity score matching for each variable included in the propensity score. A relative standardized difference less than 10% was considered to support the assumption of balance between groups.

