## supplemental tables for "Impaired renal gluconeogenesis is a major determinant of acute kidney injury associated mortality"

### Supplemental table 1: Baseline data from patients undergoing renal vein catheterization

| | Control (no AKI) n=87 | AKI n=18 | Pvalue |
|---|---|---|---|
| age (years) | 66.28 (21.18) | 68.8 (13) | 0.5679 |
| sex male | 84 (88%) | 9 (75%) | 0.9025 |
| hypertension | 58 (61%) | 9 (75%) | 0.8431 |
| beta-blockers | 78 (82%) | 10 (83%) | 1 |
| LVEF (%) | 55.6 (5.38) | 43.1 (0.63) | p<0.001 |
| serum creatinine at ICU admission (µmol/L) | 82.48 (7.16) | 90.5 (3.9) | p<0.001 |
| cardiopulmonary bypass time (min) | 76.89 (25.29) | 182.7 (23.7) | p<0.001 |
| clamping time (min) | 47.23 (15.98) | 104.3 (17) | p<0.001 |
| renal blood flow (mL/min) | 698.89 (210.68) | 459.46 (176.32) | p<0.001 |
| cardiac Index (L/min/m2) | 3.05 (0.91) | 2.63 (0.08) | p<0.001 |
| urine output (mL/min) | 3.41 (1.04) | 4.04 (0.48) | 0.0013 |
| first measured GFR (mL/min) | 64.87 (16.24) | 32.3 (3.6) | p<0.001 |
| renal O2 extraction (%) | 10.66 (2.97) | 16.3 (0.9) | p<0.001 |

### Supplemental table 2: Baseline characteristics of patients in ICU

| | Control (no AKI) (n=15657) | AKI (n=8616) | Total (n=24273) | p value |
|---|---|---|---|---|
| **sex male** | 9156.00 (58.5%) | 5447.00 (63.2%) | 14603.00 (60.2%) | < 0.001 |
| **age (years)** | | | | < 0.001 |
| mean (sd) | 59.08 (17.93) | 62.88 (16.49) | 60.43 (17.52) | |
| Median (Q1, Q3) | 61.00 (47.00, 73.00) | 66.00 (53.00, 75.00) | 63.00 (49.00, 74.00) | |
| Missing | 1.00 | 0.00 | 1.00 | |
| **BMI (kg/m$^2$)** | | | | < 0.001 |
| mean (sd) | 25.06 (6.95) | 27.75 (13.47) | 26.18 (10.27) | |
| Median (Q1, Q3) | 24.30 (21.91, 27.34) | 26.12 (23.37, 29.41) | 24.84 (22.49, 28.04) | |
| Missing | 3849.00 | 201.00 | 4050.00 | |
| **reason for ICU admission** | | | | < 0.001 |
| medical (neurological) | 1454.00 (9.3%) | 571.00 (6.6%) | 2025.00 (8.3%) | |
| medical (other) | 3899.00 (24.9%) | 2527.00 (29.3%) | 6426.00 (26.5%) | |
| medical (respiratory failure) | 2215.00 (14.1%) | 1610.00 (18.7%) | 3825.00 (15.8%) | |
| other | 2350.00 (15.0%) | 933.00 (10.8%) | 3283.00 (13.5%) | |
| surgery (neurological) | 98.00 (0.6%) | 34.00 (0.4%) | 132.00 (0.5%) | |
| surgical (cardiac) | 1198.00 (7.7%) | 1097.00 (12.7%) | 2295.00 (9.5%) | |
| surgical (elective) | 1476.00 (9.4%) | 769.00 (8.9%) | 2245.00 (9.2%) | |
| surgical (neurological) | 2198.00 (14.0%) | 600.00 (7.0%) | 2798.00 (11.5%) | |
| surgical (transplantation) | 397.00 (2.5%) | 197.00 (2.3%) | 594.00 (2.4%) | |
| surgical (urgent) | 372.00 (2.4%) | 278.00 (3.2%) | 650.00 (2.7%) | |
| **ICU length of stay (hours)** | | | | < 0.001 |
| mean (sd) | 67.95 (93.39) | 163.15 (214.36) | 101.74 (154.96) | |
| Median (Q1, Q3) | 40.00 (22.00, 73.00) | 92.00 (46.00, 190.00) | 50.00 (25.00, 112.00) | |
| Missing | 0.00 | 0.00 | 0.00 | |
| **SAPS** | | | | < 0.001 |
| mean (sd) | 38.06 (17.59) | 47.61 (18.48) | 41.45 (18.49) | |
| Median (Q1, Q3) | 36.00 (25.00, 50.00) | 46.00 (34.00, 60.00) | 40.00 (27.00, 54.00) | |
| Missing | 7.00 | 3.00 | 10.00 | |
| **apache** | | | | < 0.001 |
| mean (sd) | 19.62 (7.85) | 23.63 (8.38) | 21.07 (8.27) | |
| Median (Q1, Q3) | 19.00 (14.00, 25.00) | 23.00 (18.00, 29.00) | 21.00 (15.00, 27.00) | |
| Missing | 1202.00 | 455.00 | 1657.00 | |
| **death** | 975.00 (6.2%) | 1164.00 (13.5%) | 2139.00 (8.8%) | < 0.001 |
| **bilirubin (mmol/L)** | | | | < 0.001 |
| mean (sd) | 21.52 (49.48) | 26.71 (61.49) | 23.63 (54.73) | |
| Median (Q1, Q3) | 12.00 (8.00, 19.00) | 13.00 (8.00, 21.00) | 13.00 (8.00, 20.00) | |
| Missing | 5426.00 | 1621.00 | 7047.00 | |
| **prothrombin ratio (%)** | | | | < 0.001 |
| mean (sd) | 78.71 (18.08) | 76.93 (18.42) | 78.01 (18.23) | |
| Median (Q1, Q3) | 84.00 (70.00, 92.00) | 82.00 (68.00, 91.00) | 83.00 (69.00, 92.00) | |
| Missing | 5902.00 | 2329.00 | 8231.00 | |
| **ASAT (mmol/L)** | | | | < 0.001 |
| mean (sd) | 79.93 (239.10) | 92.45 (183.68) | 84.94 (218.68) | |
| Median (Q1, Q3) | 35.00 (22.00, 67.00) | 42.00 (25.00, 82.00) | 38.00 (23.00, 73.00) | |
| Missing | 4796.00 | 1363.00 | 6159.00 | |
| **ALAT (mmol/L)** | | | | < 0.001 |

| | | | | |
|---|---|---|---|---|
| mean (sd) | 61.16 (129.46) | 74.48 (141.00) | 66.59 (134.43) | |
| Median (Q1, Q3) | 26.00 (16.00, 53.00) | 29.50 (17.00, 61.00) | 28.00 (16.00, 56.00) | |
| Missing | 5577.00 | 1691.00 | 7268.00 | |
| **excess base (mmol/L)** | | | | < 0.001 |
| mean (sd) | -1.05 (4.48) | -0.72 (4.54) | -0.93 (4.50) | |
| Median (Q1, Q3) | -0.70 (-3.30, 1.60) | -0.20 (-3.30, 2.25) | -0.55 (-3.30, 1.80) | |
| Missing | 168.00 | 49.00 | 217.00 | |
| **hemoglobin (g/L)** | | | | < 0.001 |
| mean (sd) | 106.99 (20.46) | 101.90 (19.66) | 105.18 (20.32) | |
| Median (Q1, Q3) | 105.50 (90.50, 121.00) | 97.50 (86.00, 115.00) | 102.50 (88.50, 119.00) | |
| Missing | 2.00 | 0.00 | 2.00 | |
| **red blood cells administration** | | | | < 0.001 |
| mean (sd) | 0.32 (1.08) | 1.13 (4.71) | 0.61 (2.96) | |
| Median (Q1, Q3) | 0.00 (0.00, 0.00) | 0.00 (0.00, 1.00) | 0.00 (0.00, 0.00) | |
| Missing | 0.00 | 0.00 | 0.00 | |
| **central venous pressure (mmHg)** | | | | < 0.001 |
| mean (sd) | 10.44 (4.92) | 11.55 (4.90) | 11.08 (4.94) | |
| Median (Q1, Q3) | 10.00 (7.00, 13.00) | 11.00 (8.50, 14.00) | 11.00 (8.00, 14.00) | |
| Missing | 12926.00 | 4858.00 | 17784.00 | |
| **isoprenaline**<br>Nb of treated patients (%) | 69.00 (0.4%) | 92.00 (1.1%) | 161.00 (0.7%) | < 0.001 |
| **dobutamine**<br>Nb of treated patients (%) | 743.00 (4.7%) | 1432.00 (16.6%) | 2175.00 (9.0%) | < 0.001 |
| **epinephrine**<br>Nb of treated patients (%) | 246.00 (1.6%) | 391.00 (4.5%) | 637.00 (2.6%) | < 0.001 |
| **norepinephrine**<br>Nb of treated patients (%) | 4052.00 (25.9%) | 4713.00 (54.7%) | 8765.00 (36.1%) | < 0.001 |
| **terlipressine**<br>Nb of treated patients (%) | 65.00 (0.4%) | 137.00 (1.6%) | 202.00 (0.8%) | < 0.001 |
| **midazolam**<br>Nb of treated patients (%) | 1627.00 (10.4%) | 2493.00 (28.9%) | 4120.00 (17.0%) | < 0.001 |
| **levosimendan**<br>Nb of treated patients (%) | 34.00 (0.2%) | 119.00 (1.4%) | 153.00 (0.6%) | < 0.001 |
| **propofol**<br>Nb of treated patients (%) | 4288.00 (27.4%) | 4090.00 (47.5%) | 8378.00 (34.5%) | < 0.001 |
| **dexmedetomidine**<br>Nb of treated patients (%) | 228.00 (1.5%) | 465.00 (5.4%) | 693.00 (2.9%) | < 0.001 |
| **metformin**<br>Nb of treated patients (%) | 85.00 (0.5%) | 104.00 (1.2%) | 189.00 (0.8%) | < 0.001 |
| **aldactone**<br>Nb of treated patients (%) | 525.00 (3.4%) | 751.00 (8.7%) | 1276.00 (5.3%) | < 0.001 |
| **beta-blocker**<br>Nb of treated patients (%) | 2101.00 (13.4%) | 1889.00 (21.9%) | 3990.00 (16.4%) | < 0.001 |
| **KDIGO criteria** | | | | < 0.001 |
| KDIGO 1 | 0.00 (0.0%) | 4704.00 (54.6%) | 4704.00 (19.4%) | |
| KDIGO 2 | 0.00 (0.0%) | 3516.00 (40.8%) | 3516.00 (14.5%) | |
| KDIGO 3 | 0.00 (0.0%) | 396.00 (4.6%) | 396.00 (1.6%) | |
| no AKI | 15657.00 (100.0%) | 0.00 (0.0%) | 15657.00 (64.5%) | |
| **eGFR at ICU admission (ml/min/1.73m$^2$)** | | | | < 0.001 |
| mean (sd) | 80.40 (34.38) | 72.51 (33.99) | 77.60 (34.45) | |
| Median (Q1, Q3) | 87.26 (55.75, 105.42) | 75.70 (45.84, 97.75) | 83.72 (51.58, 103.18) | |
| Missing | 1.00 | 0.00 | 1.00 | |

| | | | | |
|---|---|---|---|---|
| **serum creatinine levels at ICU admission (mmol/L)** | | | | < 0.001 |
| mean (sd) | 108.45 (113.84) | 118.20 (110.18) | 111.91 (112.65) | |
| Median (Q1, Q3) | 75.00 (58.00, 106.00) | 84.00 (64.00, 125.00) | 78.00 (60.00, 114.00) | |
| Missing | 0.00 | 0.00 | 0.00 | |
| **maximal serum creatinine levels during ICU stay (mmol/L)** | | | | < 0.001 |
| mean (sd) | 111.63 (116.51) | 141.72 (128.11) | 122.31 (121.61) | |
| Median (Q1, Q3) | 77.00 (60.00, 110.00) | 97.00 (71.00, 158.00) | 83.00 (63.00, 125.00) | |
| Missing | 0.00 | 0.00 | 0.00 | |
| **urine output (mL/kg/h)** | | | | < 0.001 |
| mean (sd) | 1.54 (1.50) | 0.95 (0.53) | 1.30 (1.23) | |
| Median (Q1, Q3) | 1.34 (0.95, 1.85) | 0.90 (0.59, 1.25) | 1.14 (0.77, 1.60) | |
| Missing | 3689.00 | 86.00 | 3775.00 | |
| **total fluid balance during ICU stay (mL)** | | | | < 0.001 |
| mean (sd) | 1312.85 (13493.60) | 4024.75 (9618.94) | 2275.47 (12327.53) | |
| Median (Q1, Q3) | -131.83 (-2385.63, 2568.10) | 1602.56 (-531.16, 5433.50) | 437.55 (-1729.72, 3578.22) | |
| Missing | 0.00 | 0.00 | 0.00 | |
| **number of glucose and lactate measurements during ICU stay** | | | | < 0.001 |
| mean (sd) | 6.16 (4.13) | 7.19 (6.05) | 6.53 (4.92) | |
| Median (Q1, Q3) | 6.00 (3.93, 8.00) | 7.14 (5.33, 8.73) | 6.47 (4.41, 8.35) | |
| Missing | 0.00 | 0.00 | 0.00 | |
| **insulin dose (units/kg/h)** | | | | < 0.001 |
| mean (sd) | 0.01 (0.01) | 0.01 (0.14) | 0.01 (0.09) | |
| Median (Q1, Q3) | 0.00 (0.00, 0.00) | 0.00 (0.00, 0.01) | 0.00 (0.00, 0.01) | |
| Missing | 3689.00 | 86.00 | 3775.00 | |
| **energy intake (kcal/kg/day)** | | | | < 0.001 |
| mean (sd) | 14.27 (27.21) | 25.46 (30.90) | 18.92 (29.33) | |
| Median (Q1, Q3) | 0.95 (0.00, 14.70) | 10.89 (0.49, 45.19) | 2.82 (0.00, 29.53) | |
| Missing | 3689.00 | 86.00 | 3775.00 | |
| **glucose-Lactate group** | | | | < 0.001 |
| | 15.00 (0.1%) | 4.00 (0.0%) | 19.00 (0.1%) | |
| Baseline | 3685.00 (23.5%) | 1956.00 (22.7%) | 5641.00 (23.2%) | |
| impaired metabolism | 6149.00 (39.3%) | 3762.00 (43.7%) | 9911.00 (40.8%) | |
| isolated hyperglycemia | 524.00 (3.3%) | 349.00 (4.1%) | 873.00 (3.6%) | |
| isolated hypoglycemia | 3396.00 (21.7%) | 1154.00 (13.4%) | 4550.00 (18.7%) | |
| stress response | 1888.00 (12.1%) | 1391.00 (16.1%) | 3279.00 (13.5%) | |
| **number (%) of glucose-lactate measurements in impaired metabolism group** | | | | < 0.001 |
| mean (sd) | 35.57 (32.98) | 37.53 (27.99) | 36.27 (31.31) | |
| Median (Q1, Q3) | 29.00 (2.00, 59.00) | 33.00 (14.00, 57.00) | 31.00 (9.00, 57.00) | |
| Missing | 0.00 | 0.00 | 0.00 | |

**Supplemental table 3: Baseline characteristics of ICU patients after PSM**

| | AKI (N=8575) | no AKI (N=17150) | Total (N=25725) | Standardized difference |
|---|---|---|---|---|
| **SAPS** | | | | 4.839 |
| mean (sd) | 47.25 (18.56) | 48.15 (19.60) | 47.85 (19.26) | |
| Median (Q1, Q3) | 46.00 (33.00, 60.00) | 47.00 (34.00, 61.00) | 47.00 (34.00, 61.00) | |
| Missing | 0.00 | 0.00 | 0.00 | |
| **eGFR at ICU admission** | | | | 3.735 |
| mean (sd) | 82.69 (50.89) | 80.79 (50.62) | 81.42 (50.72) | |
| Median (Q1, Q3) | 77.00 (48.00, 107.00) | 77.00 (43.00, 109.00) | 77.00 (45.00, 109.00) | |
| Missing | 0.00 | 0.00 | 0.00 | |
| **sex male** | 5426.00 (63.3%) | 10928.00 (63.7%) | 16354.00 (63.6%) | 0.919 |
| **age (years)** | | | | 2.277 |
| mean (sd) | 62.87 (16.50) | 63.25 (16.52) | 63.12 (16.52) | |
| Median (Q1, Q3) | 66.00 (53.00, 75.00) | 66.00 (53.00, 76.00) | 66.00 (53.00, 76.00) | |
| Missing | 0.00 | 0.00 | 0.00 | |
| **BMI (kg/m$^2$)** | | | | 2.308 |
| mean (sd) | 27.66 (13.28) | 27.35 (11.16) | 27.45 (11.91) | |
| Median (Q1, Q3) | 25.98 (23.37, 29.39) | 25.31 (23.84, 28.31) | 25.31 (23.55, 29.05) | |
| Missing | 0.00 | 0.00 | 0.00 | |
| **bilirubin (mmol/L)** | | | | 1.266 |
| mean (sd) | 26.82 (57.55) | 27.55 (61.53) | 27.30 (60.23) | |
| Median (Q1, Q3) | 16.00 (10.00, 21.00) | 19.09 (10.00, 20.00) | 18.00 (10.00, 20.28) | |
| Missing | 0.00 | 0.00 | 0.00 | |
| **prothrombin ratio** | | | | 5.170 |
| mean (sd) | 75.75 (16.46) | 74.89 (17.38) | 75.18 (17.08) | |
| Median (Q1, Q3) | 78.12 (69.00, 86.00) | 78.00 (69.00, 85.00) | 78.03 (69.00, 86.00) | |
| Missing | 0.00 | 0.00 | 0.00 | |
| **hemoglobin (g/L)** | | | | 1.410 |
| mean (sd) | 102.32 (19.96) | 102.04 (19.74) | 102.14 (19.81) | |
| Median (Q1, Q3) | 98.00 (86.50, 116.00) | 99.00 (86.50, 115.00) | 99.00 (86.50, 115.00) | |
| Missing | 0.00 | 0.00 | 0.00 | |
| **insulin dose (unit/kg/hour)** | | | | 0.670 |
| mean (sd) | 0.01 (0.01) | 0.01 (0.02) | 0.01 (0.02) | |
| Median (Q1, Q3) | 0.00 (0.00, 0.01) | 0.00 (0.00, 0.01) | 0.00 (0.00, 0.01) | |
| Missing | 0.00 | 0.00 | 0.00 | |
| **energy intake (kcal/kg/day)** | | | | 4.581 |
| mean (sd) | 25.91 (36.70) | 24.23 (34.41) | 24.79 (35.20) | |
| Median (Q1, Q3) | 8.39 (0.29, 39.80) | 13.48 (0.26, 34.36) | 12.99 (0.27, 36.61) | |
| Missing | 0.00 | 0.00 | 0.00 | |
| **dobutamine** | 1007.00 (11.7%) | 1866.00 (10.9%) | 2873.00 (11.2%) | 2.680 |
| **epinephrine** | 166.00 (1.9%) | 310.00 (1.8%) | 476.00 (1.9%) | 0.931 |
| **norepinephrine** | 2532.00 (29.5%) | 4860.00 (28.3%) | 7392.00 (28.7%) | 2.607 |
| **midazolam** | 1420.00 (16.6%) | 2657.00 (15.5%) | 4077.00 (15.8%) | 2.870 |
| **propofol** | 2219.00 (25.9%) | 4359.00 (25.4%) | 6578.00 (25.6%) | 1.052 |
| **metformin** | 39.00 (0.5%) | 66.00 (0.4%) | 105.00 (0.4%) | 1.040 |
| **beta blocker** | 1684.00 (19.6%) | 3184.00 (18.6%) | 4868.00 (18.9%) | 2.701 |
| **terlipressine** | 46.00 (0.5%) | 87.00 (0.5%) | 133.00 (0.5%) | 0.399 |

**Supplemental table 4: Baseline characteristics of allograft patients**

| | Living donor (N=153) | Deceased donor (N=122) | Total (N=275) | p value |
|---|---|---|---|---|
| **sex male** | 101.00 (66.0%) | 75.00 (61.5%) | 176.00 (64.0%) | 0.447 |
| **Age (years)** | | | | 0.043 |
| meansd | 52.12 (13.99) | 55.48 (13.66) | 53.61 (13.92) | |
| Median (Q1, Q3) | 52.00 (42.00, 63.00) | 57.00 (45.25, 67.75) | 54.00 (43.00, 65.00) | |
| Missing | 0.00 | 0.00 | 0.00 | |
| **BMI (kg/m$^2$)** | | | | 0.158 |
| meansd | 26.99 (21.89) | 26.03 (4.68) | 26.56 (16.59) | |
| Median (Q1, Q3) | 24.41 (22.41, 27.78) | 25.67 (22.88, 29.37) | 25.11 (22.49, 28.63) | |
| Missing | 23.00 | 18.00 | 41.00 | |
| **ICU length of stay (hours)** | | | | 0.049 |
| meansd | 36.49 (22.20) | 42.25 (26.36) | 39.04 (24.26) | |
| Median (Q1, Q3) | 25.00 (22.00, 45.00) | 36.00 (23.25, 50.00) | 33.00 (22.00, 46.00) | |
| Missing | 0.00 | 0.00 | 0.00 | |
| **SAPS** | | | | 0.041 |
| meansd | 26.06 (13.17) | 29.07 (12.69) | 27.40 (13.02) | |
| Median (Q1, Q3) | 24.00 (18.00, 32.00) | 27.00 (20.00, 36.00) | 25.00 (18.00, 34.00) | |
| Missing | 0.00 | 0.00 | 0.00 | |
| **apache** | | | | 0.513 |
| meansd | 18.38 (5.91) | 18.88 (6.15) | 18.60 (6.01) | |
| Median (Q1, Q3) | 18.00 (15.00, 21.50) | 18.00 (15.00, 22.00) | 18.00 (15.00, 22.00) | |
| Missing | 10.00 | 10.00 | 20.00 | |
| **death** | 0.00 (0.0%) | 0.00 (0.0%) | 0.00 (0.0%) | 1.000 |
| **bilirubin (mmol/L)** | | | | 0.053 |
| meansd | 8.82 (5.46) | 10.22 (5.64) | 9.39 (5.56) | |
| Median (Q1, Q3) | 7.00 (5.00, 12.00) | 10.00 (6.00, 13.00) | 8.00 (5.00, 12.00) | |
| Missing | 33.00 | 39.00 | 72.00 | |
| **prothrombin ratio (%)** | | | | 0.227 |
| meansd | 86.37 (9.03) | 87.59 (10.22) | 86.95 (9.60) | |
| Median (Q1, Q3) | 87.75 (79.75, 94.00) | 90.00 (82.50, 96.00) | 89.00 (80.50, 95.00) | |
| Missing | 69.00 | 47.00 | 116.00 | |
| **ASAT (mmol/L)** | | | | 0.929 |
| meansd | 34.69 (34.04) | 39.02 (42.95) | 36.56 (38.13) | |
| Median (Q1, Q3) | 27.00 (21.00, 39.50) | 26.50 (21.00, 40.25) | 27.00 (21.00, 40.00) | |
| Missing | 30.00 | 28.00 | 58.00 | |
| **ALAT (mmol/L)** | | | | 0.202 |
| meansd | 33.25 (37.97) | 33.26 (47.11) | 33.26 (42.05) | |
| Median (Q1, Q3) | 23.00 (16.00, 36.25) | 19.00 (15.50, 32.50) | 22.00 (16.00, 35.00) | |
| Missing | 33.00 | 31.00 | 64.00 | |
| **excess base (mmol/L)** | | | | 0.785 |
| meansd | -3.01 (3.81) | -3.20 (3.40) | -3.09 (3.63) | |
| Median (Q1, Q3) | -3.00 (-5.40, -0.60) | -3.00 (-5.75, -0.55) | -3.00 (-5.61, -0.60) | |
| Missing | 4.00 | 3.00 | 7.00 | |
| **hemoglobin (g/L)** | | | | 0.882 |
| meansd | 96.05 (13.18) | 95.66 (12.98) | 95.88 (13.07) | |
| Median (Q1, Q3) | 96.00 (86.00, 104.00) | 95.25 (87.00, 103.00) | 95.50 (86.25, 104.00) | |
| Missing | 0.00 | 0.00 | 0.00 | |

| | | | | |
|---|---|---|---|---|
| **red blood cells administration** | | | | 0.195 |
| meansd | 0.05 (0.29) | 0.13 (0.60) | 0.08 (0.46) | |
| Median (Q1, Q3) | 0.00 (0.00, 0.00) | 0.00 (0.00, 0.00) | 0.00 (0.00, 0.00) | |
| Missing | 0.00 | 0.00 | 0.00 | |
| **isoprenaline** Nb of treated patients (%) | 0.00 (0.0%) | 0.00 (0.0%) | 0.00 (0.0%) | 1.000 |
| **dobutamine** Nb of treated patients (%) | 8.00 (5.2%) | 9.00 (7.4%) | 17.00 (6.2%) | 0.601 |
| **epinephrine** Nb of treated patients (%) | 1.00 (0.7%) | 4.00 (3.3%) | 5.00 (1.8%) | 0.176 |
| **norepinephrine** Nb of treated patients (%) | 40.00 (26.1%) | 38.00 (31.1%) | 78.00 (28.4%) | 0.418 |
| **terlipressine** Nb of treated patients (%) | 1.00 (0.7%) | 0.00 (0.0%) | 1.00 (0.4%) | 1.000 |
| **midazolam** Nb of treated patients (%) | 14.00 (9.2%) | 14.00 (11.5%) | 28.00 (10.2%) | 0.545 |
| **levosimendan** Nb of treated patients (%) | 0.00 (0.0%) | 0.00 (0.0%) | 0.00 (0.0%) | 1.000 |
| **Propofol** Nb of treated patients (%) | 35.00 (22.9%) | 26.00 (21.3%) | 61.00 (22.2%) | 0.790 |
| **Dexmedetomidine** Nb of treated patients (%) | 0.00 (0.0%) | 0.00 (0.0%) | 0.00 (0.0%) | 1.000 |
| **Metformine** Nb of treated patients (%) | 0.00 (0.0%) | 0.00 (0.0%) | 0.00 (0.0%) | 1.000 |
| **Aldactone** Nb of treated patients (%) | 2.00 (1.3%) | 3.00 (2.5%) | 5.00 (1.8%) | 0.651 |
| **beta-blocker** Nb of treated patients (%) | 27.00 (17.6%) | 18.00 (14.8%) | 45.00 (16.4%) | 0.629 |
| **serum creatinine levels at ICU admission (mmol/L)** | | | | 0.591 |
| meansd | 473.35 (190.72) | 482.43 (186.09) | 477.38 (188.39) | |
| Median (Q1, Q3) | 451.00 (352.00, 567.00) | 464.50 (360.25, 593.50) | 454.00 (356.50, 573.00) | |
| Missing | 0.00 | 0.00 | 0.00 | |
| **maximal serum creatinine levels during ICU stay (mmol/L)** | | | | 0.341 |
| meansd | 488.33 (195.59) | 508.30 (199.63) | 497.19 (197.28) | |
| Median (Q1, Q3) | 467.00 (365.00, 595.00) | 486.50 (366.25, 626.75) | 473.00 (365.50, 615.50) | |
| Missing | 0.00 | 0.00 | 0.00 | |
| **urine output (mL/kg/h)** | | | | < 0.001 |
| meansd | 3.96 (2.44) | 2.73 (2.22) | 3.41 (2.42) | |
| Median (Q1, Q3) | 3.95 (1.97, 5.24) | 2.23 (0.99, 3.59) | 2.97 (1.41, 4.81) | |
| Missing | 22.00 | 18.00 | 40.00 | |
| **total fluid balance during ICU stay (mL)** | | | | < 0.001 |
| meansd | 1882.81 (5556.48) | 4425.28 (5984.34) | 3010.75 (5877.39) | |
| Median (Q1, Q3) | 1303.19 (-1487.61, 4727.26) | 3234.64 (1109.74, 7260.88) | 2308.79 (-386.87, 5698.56) | |
| Missing | 0.00 | 0.00 | 0.00 | |
| **number of glucose and lactate measurements during ICU stay** | | | | 0.130 |
| meansd | 6.21 (1.94) | 6.71 (2.34) | 6.43 (2.14) | |
| Median (Q1, Q3) | 6.26 (5.05, 7.43) | 6.42 (5.25, 8.00) | 6.26 (5.17, 7.65) | |
| Missing | 0.00 | 0.00 | 0.00 | |
| **insulin dose (units/kg/h)** | | | | 0.081 |
| meansd | 0.01 (0.02) | 0.01 (0.01) | 0.01 (0.01) | |

| | | | | |
|---|---|---|---|---|
| Median (Q1, Q3) | 0.00 (0.00, 0.01) | 0.01 (0.00, 0.01) | 0.00 (0.00, 0.01) | |
| Missing | 22.00 | 18.00 | 40.00 | |
| **energy intake (kcal/kg/day)** | | | | 0.013 |
| meansd | 2.03 (9.50) | 4.97 (12.33) | 3.33 (10.92) | |
| Median (Q1, Q3) | 0.00 (0.00, 0.00) | 0.00 (0.00, 1.02) | 0.00 (0.00, 0.14) | |
| Missing | 22.00 | 18.00 | 40.00 | |
| **Serum creatinine levels at 6 months (mmol/L)** | | | | 0.406 |
| meansd | 130.18 (54.96) | 178.12 (236.49) | 151.29 (163.59) | |
| Median (Q1, Q3) | 118.00 (102.00, 146.50) | 132.50 (93.25, 159.50) | 122.00 (99.00, 152.00) | |
| Missing | 3.00 | 4.00 | 7.00 | |
| **maximal serum creatinine levels at 15 days stay (mmol/L)** | | | | < 0.001 |
| meansd | 152.09 (117.22) | 212.00 (193.26) | 178.67 (158.13) | |
| Median (Q1, Q3) | 121.00 (98.00, 154.00) | 145.50 (111.25, 211.00) | 131.00 (103.00, 173.00) | |
| Missing | 0.00 | 0.00 | 0.00 | |
| **number (%) of glucose-lactate measurements in impaired metabolism group** | | | | 0.025 |
| meansd | 26.24 (24.16) | 32.76 (25.52) | 29.13 (24.94) | |
| Median (Q1, Q3) | 23.00 (0.00, 40.00) | 27.00 (14.25, 50.00) | 25.00 (9.50, 44.00) | |
| Missing | 0.00 | 0.00 | 0.00 | |
| **glucose-Lactate group** | | | | 0.253 |
| | 1.00 (0.7%) | 0.00 (0.0%) | 1.00 (0.4%) | |
| Baseline | 42.00 (27.5%) | 29.00 (23.8%) | 71.00 (25.8%) | |
| impaired metabolism | 39.00 (25.5%) | 45.00 (36.9%) | 84.00 (30.5%) | |
| isolated hyperglycemia | 16.00 (10.5%) | 7.00 (5.7%) | 23.00 (8.4%) | |
| isolated hypoglycemia | 6.00 (3.9%) | 3.00 (2.5%) | 9.00 (3.3%) | |
| stress response | 49.00 (32.0%) | 38.00 (31.1%) | 87.00 (31.6%) | |

**Supplemental table 5: Baseline characteristics of ICU AKI patients receiving or not thiamine supplementation after PSM**

| | No thiamine (N=694) | Thiamine (N=347) | Total (N=1041) | Standardized difference |
|---|---|---|---|---|
| **SAPS** | | | | 0.72 |
| mean (sd) | 55.22 (18.00) | 55.36 (18.74) | 55.27 (18.25) | |
| Median (Q1, Q3) | 56.00 (43.25, 68.00) | 56.00 (42.00, 68.00) | 56.00 (43.00, 68.00) | |
| Missing | 0.00 | 0.00 | 0.00 | |
| **eGFR at ICU admission** | | | | 5.59 |
| mean (sd) | 78.79 (36.11) | 76.81 (35.41) | 78.13 (35.88) | |
| Median (Q1, Q3) | 85.82 (51.38, 105.31) | 80.33 (51.84, 102.30) | 85.15 (51.72, 104.66) | |
| Missing | 0.00 | 0.00 | 0.00 | |
| **sex male** | 506.00 (72.9%) | 255.00 (73.5%) | 761.00 (73.1%) | 1.3 |
| **age (years)** | | | | 9.08 |
| mean (sd) | 58.83 (17.66) | 60.21 (15.27) | 59.29 (16.90) | |
| Median (Q1, Q3) | 62.00 (48.25, 72.00) | 62.00 (51.00, 72.00) | 62.00 (50.00, 72.00) | |
| Missing | 0.00 | 0.00 | 0.00 | |
| **BMI (kg/m$^2$)** | | | | 0.93 |
| mean (sd) | 26.46 (5.75) | 26.51 (6.07) | 26.48 (5.86) | |
| Median (Q1, Q3) | 24.85 (23.15, 28.40) | 25.71 (23.09, 28.94) | 25.22 (23.15, 28.58) | |
| Missing | 0.00 | 0.00 | 0.00 | |
| **bilirubin (mmol/L)** | | | | 8.51 |
| mean (sd) | 25.58 (61.77) | 31.38 (68.14) | 27.51 (63.99) | |
| Median (Q1, Q3) | 10.00 (6.00, 19.00) | 12.00 (7.00, 24.50) | 10.00 (6.00, 20.00) | |
| Missing | 0.00 | 0.00 | 0.00 | |
| **prothrombin ratio** | | | | 7.37 |
| mean (sd) | 74.63 (18.09) | 73.20 (19.44) | 74.16 (18.55) | |
| Median (Q1, Q3) | 79.00 (67.00, 88.00) | 78.00 (63.00, 88.00) | 78.49 (66.00, 88.00) | |
| Missing | 0.00 | 0.00 | 0.00 | |
| **hemoglobin (g/L)** | | | | 1.33 |
| mean (sd) | 101.59 (21.76) | 101.31 (20.86) | 101.50 (21.45) | |
| Median (Q1, Q3) | 96.50 (84.50, 117.25) | 96.00 (85.00, 116.00) | 96.00 (84.50, 116.00) | |
| Missing | 0.00 | 0.00 | 0.00 | |
| **insulin dose (unit/kg/hour)** | | | | 0.16 |
| mean (sd) | 0.01 (0.02) | 0.01 (0.02) | 0.01 (0.02) | |
| Median (Q1, Q3) | 0.00 (0.00, 0.01) | 0.00 (0.00, 0.01) | 0.00 (0.00, 0.01) | |
| Missing | 0.00 | 0.00 | 0.00 | |
| **energy intake (kcal/kg/day)** | | | | 3.3 |
| mean (sd) | 35.26 (35.44) | 34.29 (29.46) | 34.94 (33.55) | |
| Median (Q1, Q3) | 26.93 (2.51, 55.57) | 30.25 (6.76, 53.47) | 28.10 (3.05, 54.29) | |
| Missing | 0.00 | 0.00 | 0.00 | |
| **dobutamine** | 102.00 (14.7%) | 56.00 (16.1%) | 158.00 (15.2%) | 3.91 |
| **epinephrine** | 24.00 (3.5%) | 14.00 (4.0%) | 38.00 (3.7%) | 2.92 |
| **norepinephrine** | 498.00 (71.8%) | 253.00 (72.9%) | 751.00 (72.1%) | 2.59 |
| **midazolam** | 319.00 (46.0%) | 162.00 (46.7%) | 481.00 (46.2%) | 1.44 |
| **propofol** | 505.00 (72.8%) | 243.00 (70.0%) | 748.00 (71.9%) | 5.97 |
| **metformin** | 8.00 (1.2%) | 5.00 (1.4%) | 13.00 (1.2%) | 2.41 |
| **beta blocker** | 145.00 (20.9%) | 69.00 (19.9%) | 214.00 (20.6%) | 2.52 |
| **terlipressine** | 38.00 (5.5%) | 25.00 (7.2%) | 63.00 (6.1%) | 6.68 |

**Supplemental table 6: Baseline characteristics of ICU non-AKI patients receiving or not thiamine supplementation after PSM**

| | No thiamine (N=526) | Thiamine (N=263) | Total (N=789) | Standardized difference |
|---|---|---|---|---|
| **SAPS** | | | | 0.41 |
| mean (sd) | 48.34 (18.29) | 48.27 (18.70) | 48.32 (18.42) | |
| Median (Q1, Q3) | 49.00 (36.00, 61.00) | 48.00 (34.50, 61.00) | 49.00 (35.00, 61.00) | |
| Missing | 0.00 | 0.00 | 0.00 | |
| **eGFR at ICU admission** | | | | 8.31 |
| mean (sd) | 78.39 (35.67) | 75.36 (36.48) | 77.38 (35.94) | |
| Median (Q1, Q3) | 83.84 (53.26, 104.76) | 81.87 (46.92, 103.94) | 83.71 (50.95, 104.09) | |
| Missing | 0.00 | 0.00 | 0.00 | |
| **sex male** | 424.00 (80.6%) | 200.00 (76.0%) | 624.00 (79.1%) | 10.67 |
| **age (years)** | | | | 2.33 |
| mean (sd) | 56.53 (18.34) | 56.85 (13.56) | 56.64 (16.89) | |
| Median (Q1, Q3) | 59.50 (45.00, 71.00) | 58.00 (48.00, 66.00) | 58.00 (46.00, 68.00) | |
| Missing | 0.00 | 0.00 | 0.00 | |
| **BMI (kg/m$^2$)** | | | | 1.3 |
| mean (sd) | 24.81 (4.65) | 24.88 (4.86) | 24.84 (4.72) | |
| Median (Q1, Q3) | 24.30 (22.33, 26.33) | 24.66 (21.89, 27.68) | 24.49 (22.09, 26.64) | |
| Missing | 0.00 | 0.00 | 0.00 | |
| **bilirubin (mmol/L)** | | | | 8.18 |
| mean (sd) | 20.85 (41.34) | 26.02 (63.22) | 22.57 (49.73) | |
| Median (Q1, Q3) | 10.00 (6.00, 18.00) | 10.00 (6.00, 18.50) | 10.00 (6.00, 18.00) | |
| Missing | 0.00 | 0.00 | 0.00 | |
| **prothrombin ratio** | | | | 7.23 |
| mean (sd) | 76.75 (15.94) | 75.51 (17.13) | 76.34 (16.34) | |
| Median (Q1, Q3) | 79.00 (70.00, 88.00) | 78.92 (67.50, 86.75) | 79.00 (69.00, 88.00) | |
| Missing | 0.00 | 0.00 | 0.00 | |
| **hemoglobin (g/L)** | | | | 5.96 |
| mean (sd) | 110.53 (22.30) | 109.18 (22.65) | 110.08 (22.41) | |
| Median (Q1, Q3) | 111.00 (90.00, 128.00) | 108.00 (90.75, 125.00) | 110.00 (90.00, 126.00) | |
| Missing | 0.00 | 0.00 | 0.00 | |
| **insulin dose (unit/kg/hour)** | | | | 4.4 |
| mean (sd) | 0.01 (0.02) | 0.01 (0.02) | 0.01 (0.02) | |
| Median (Q1, Q3) | 0.00 (0.00, 0.01) | 0.00 (0.00, 0.00) | 0.00 (0.00, 0.00) | |
| Missing | 0.00 | 0.00 | 0.00 | |
| **energy intake (kcal/kg/day)** | | | | 0.15 |
| mean (sd) | 22.05 (32.53) | 22.09 (26.84) | 22.06 (30.73) | |
| Median (Q1, Q3) | 4.60 (0.29, 36.68) | 9.32 (1.45, 35.56) | 6.06 (0.51, 36.51) | |
| Missing | 0.00 | 0.00 | 0.00 | |
| **dobutamine** | 30.00 (5.7%) | 18.00 (6.8%) | 48.00 (6.1%) | 4.51 |
| **epinephrine** | 8.00 (1.5%) | 2.00 (0.8%) | 10.00 (1.3%) | 8.74 |
| **norepinephrine** | 212.00 (40.3%) | 117.00 (44.5%) | 329.00 (41.7%) | 8.4 |
| **midazolam** | 128.00 (24.3%) | 67.00 (25.5%) | 195.00 (24.7%) | 2.61 |
| **propofol** | 276.00 (52.5%) | 132.00 (50.2%) | 408.00 (51.7%) | 4.55 |
| **metformin** | 1.00 (0.2%) | 1.00 (0.4%) | 2.00 (0.3%) | 3.08 |
| **beta blocker** | 54.00 (10.3%) | 29.00 (11.0%) | 83.00 (10.5%) | 2.42 |
| **terlipressine** | 1.00 (0.2%) | 1.00 (0.4%) | 2.00 (0.3%) | 3.08 |
