## extended data for "Impaired renal gluconeogenesis is a major determinant of acute kidney injury associated mortality"

### EXTENDED DATA FIGURES

EXTENDED DATA FIGURE 1: VALIDATION OF THE NEPH-GFP MOUSE 2

EXTENDED DATA FIGURE 2: A MOUSE MODEL OF MODERATE IRI TO STUDY TUBULAR REPAIR 3

EXTENDED DATA FIGURE 3: NUCSEQ ANALYSIS AFTER IRI 4

EXTENDED DATA FIGURE 4 MRNA EXPRESSION OF GLUCONEOGENESIS REGULATORS 5

EXTENDED DATA FIGURE 5: GLUCONEOGENESIS AND GLYCOLYSIS GENES IN HUMAN KIDNEY BIOPSIES CLASSIFIED USING A COMPUTATIONAL APPROACH. 6

EXTENDED DATA FIGURE 6: FBP1 AND PKM EXPRESSION ACCORDING TO ONE-YEAR EGFR 7

EXTENDED DATA FIGURE 7: GLUCOSE AND LACTATE PATTERNS DEFINITION 8

EXTENDED DATA FIGURE 8: BALANCE OF VARIABLES FOR AKI BEFORE AND AFTER MATCHING 9

EXTENDED DATA FIGURE 9: FLOW CHART AFTER PSM MATCHING AFTER MATCHING IN THE AKI DATASET 10

EXTENDED DATA FIGURE 10: RELATION BETWEEN IMPAIRED METABOLISM PATTERN RATE FOR EACH KIDNEY ALLOGRAFT RECIPIENT AND POD15 SERUM CREATININE. 11

EXTENDED DATA FIGURE 11: ICU MORTALITY ACCORDING TO KDIGO AND IMPAIRED METABOLISM STATUS 12

EXTENDED DATA FIGURE 12: FLOW CHART AFTER MATCHING IN THE THIAMINE DATASET 13

EXTENDED DATA FIGURE 13: BALANCE OF VARIABLES BEFORE AND AFTER MATCHING IN THE THIAMINE DATASET 14

Extended data figure 1: validation of the Neph-GFP mouse

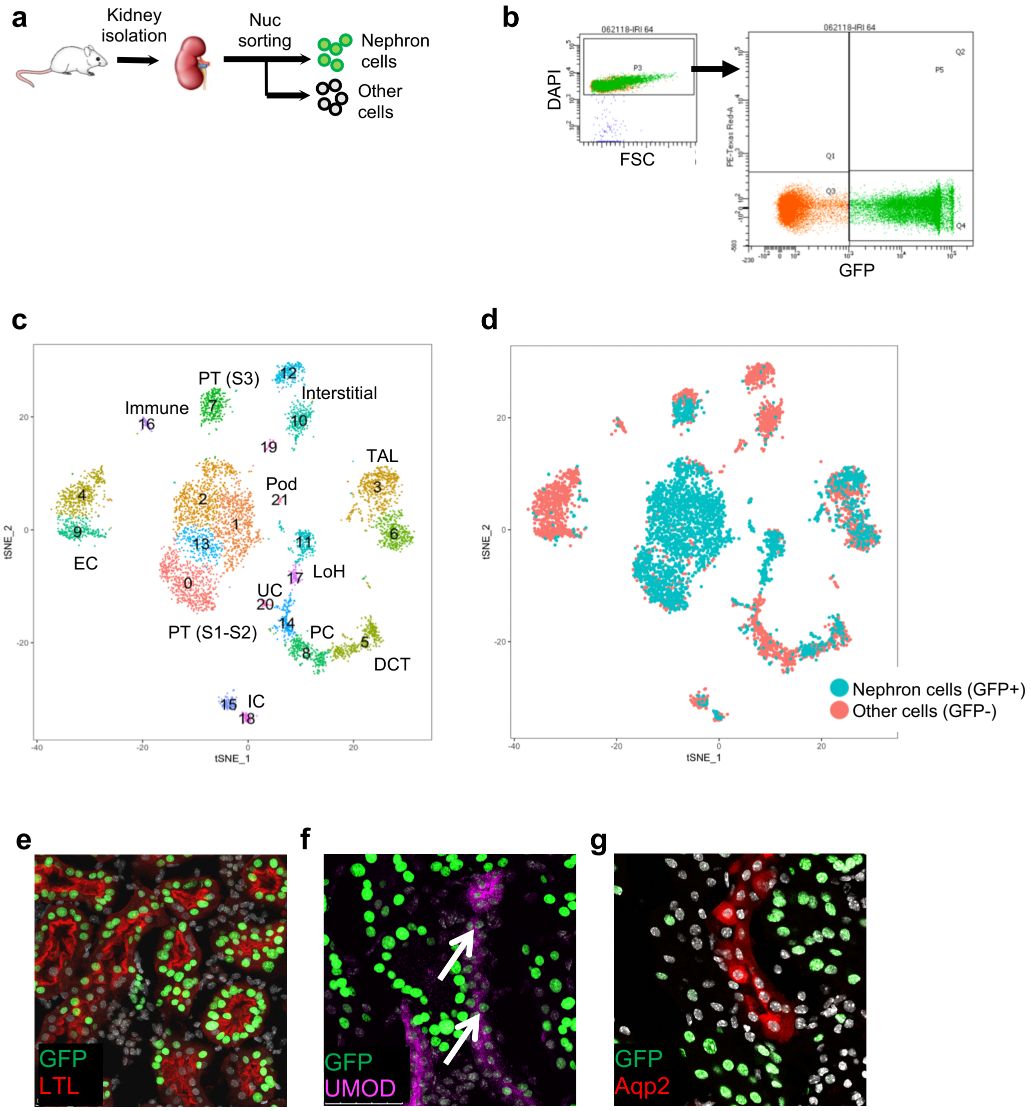


Validation of the Neph-GFP mouse. (a) Schematic representation of the separation in nephron and non-nephron cells in Neph-GFP mice. (b) Representative example of flow cytometry and sorting of nuclei isolated from the kidney of a Neph-GFP mouse. (c-d) TSNE analysis of merged GFP$^+$ and GFP$^-$ cells from one representative Neph-GFP kidney. Cell clusters are labelled with numbers and cell populations with initials in (c) (s. Fig. 2d), the separation in GFP$^+$ and GFP$^-$ cells is shown in (d). Please note that a subset of nephron cells was detected in the GFP$^-$ fraction, probably as a result of the delay in the replacement of histone proteins. The presence of cells with a transcriptional profile overlapping with principal and intercalated cells of the collecting duct in the GFP$^+$ fraction was consistent with previous observations. (e) Dotplots of representative marker genes for each cell cluster. Cell clusters are labelled with numbers, cell populations with initials, as follows: Pod: podocytes, UC: urothelial cells, IC: intercalated cells of the collecting duct, PC: principal cells of the collecting duct, PT: proximal tubule, S1-S2: segment 1 and 2, S3: segment 3, DTL: descending thin limb, LoH: Loop of Henle, EC: endothelial cells, TAL: Thick ascending limb, DCT: distal convoluted tubule. (f-h) Immunofluorescence from a representative example of a Neph-GFP mouse kidney showing GFP$^+$ nuclei in Ltl$^+$ PT cells and Umod$^+$ TAL cells, but GFP$^-$ nuclei in Aqp2$^+$ collecting duct cells.

Extended data figure 2: a mouse model of moderate IRI to study tubular repair

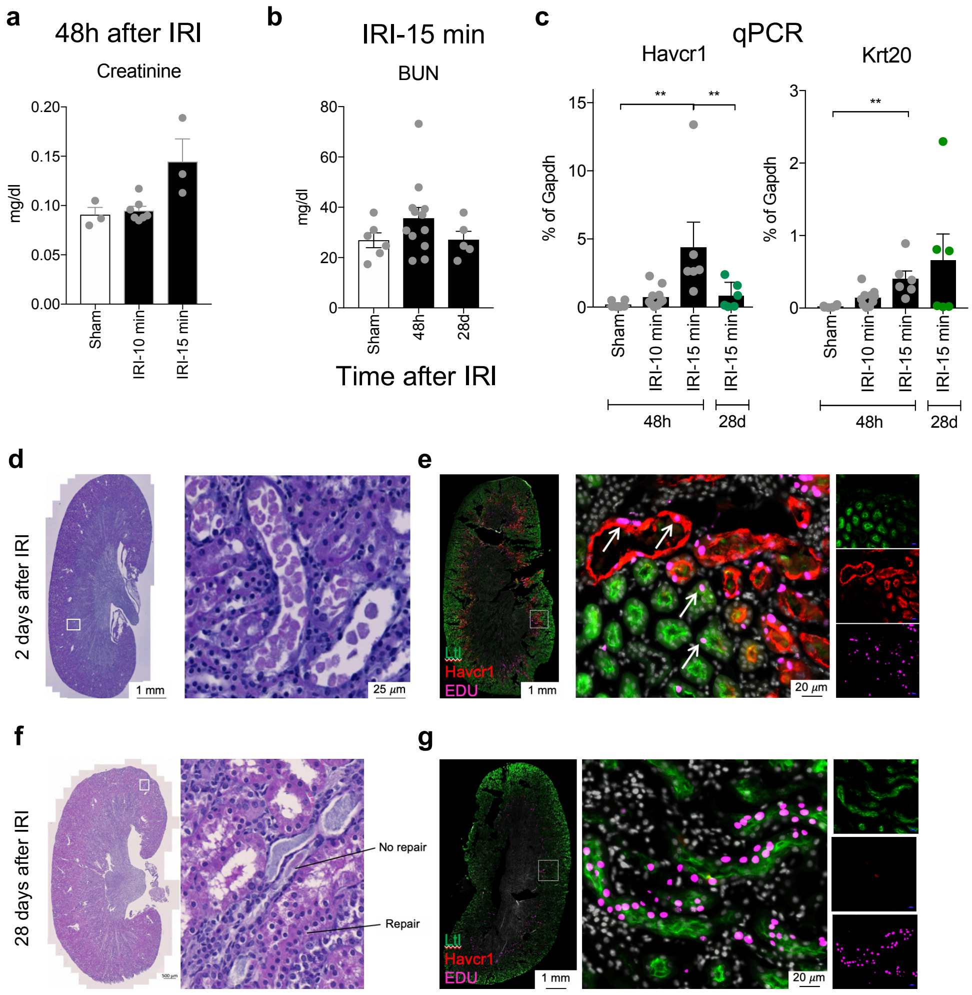


A mouse model of moderate IRI to study tubular repair. (a) Serum creatinine values measured 48h after IRI, with different ischemia times, as indicated. N=3-6 mice. (b) BUN values measure 48h and 28 days after IRI (15 minutes ischemia). N=5-12 mice. (c) qPCR on renal tissue isolated at different time points after IRI with different ischemia times, as indicated. N=5-8 mice, **P<0.01. (d-g) Conventional H&E staining (d, f) and immunofluorescence (e, g) on kidney sections obtained 2 and 28 days after 15 min IRI showing the presence of damaged tubular cell at the cortico-medullary junction (S3 region of the PT). 28 days after IRI most tubular cells recovered, but still retained EDU (injected 48h after IRI), but some flattened cells and damaged tubules persisted (f).

### Extended data figure 3: NucSeq analysis after IRI

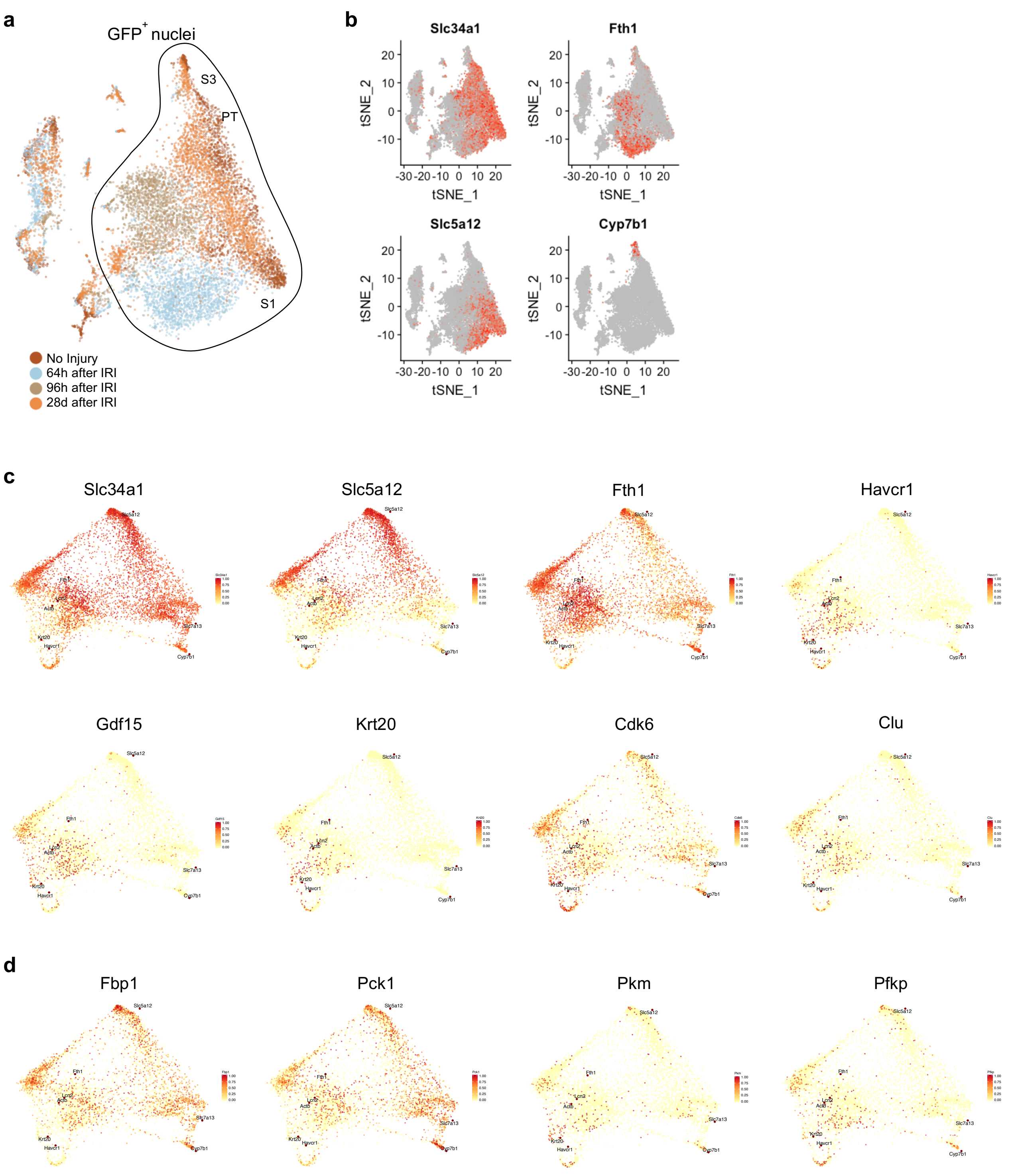


NucSeq analysis after IRI. (a) TSNE analysis on merged datasets including GFP+ nuclei at different time points after IRI as indicated. The line delineates proximal tubule (PT) cells. S1 and S3 indicate the corresponding segments of the PT. (b) Feature plots of representative genes, typically expressed by proximal tubule cells (*Slc34a1*), S1 cells (*Slc5a12*), S3 cells (*Cyp7b1*) and injured proximal tubule cells (*Fth1*). (c) SWNE plot including PT nuclei isolated at different time points after IRI as presented in Fig. 2. The colors indicate the expression level of representative genes as indicated.

Extended data figure 4 mRNA expression of gluconeogenesis regulators

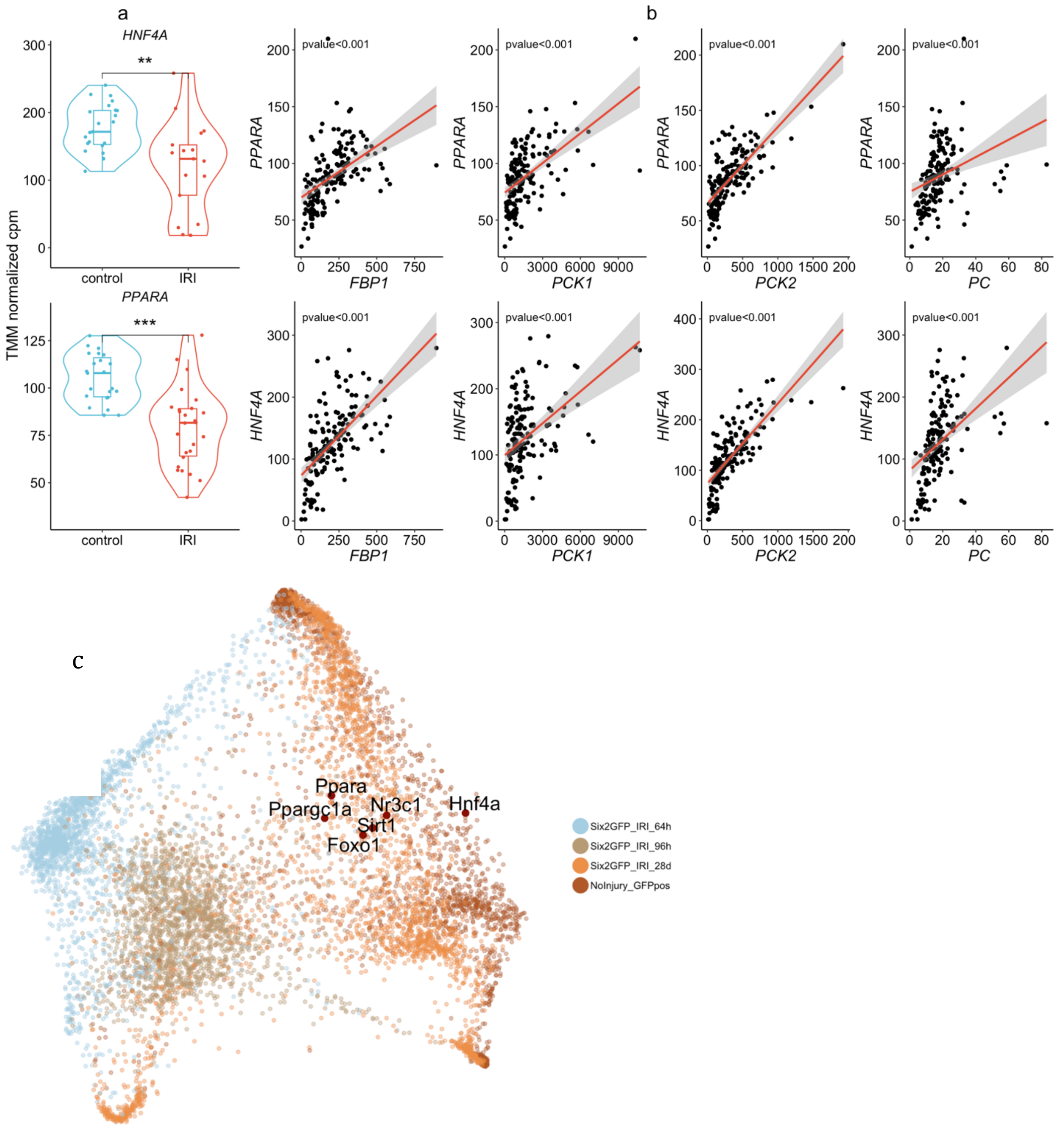


mRNA expression of gluconeogenesis regulators, *HNFA4* and *PPARA*. (a) TMM normalised cpm of HNFA4 and PPARA genes in allograft kidney biopsies obtained at the end of the transplantation from brain-stem death donor (IRI group) and 3 or 12 months after transplantation in patients with recovery renal graft status (control group). (b) relation between *HNFA4* and *PPARA* genes with gluconeogenesis genes *FBP1*, *PCK1 PCK2* and *PC.* (c) SWNE plots including PT nuclei isolated at different time points after IRI and highlighting the separation of PT cells in the early phase after IRI from steady state conditions. Representative regulators genes expressed in normal PT cells are embedded The color of the dots corresponds to the time point of harvesting after IRI. Violin plot and boxplot (with mean, IQR, 1$^{st}$ and 95$^{th}$ percentiles) are shown. Differential expression analyses were performed using genewise negative binomial model with quasi-likelihood test. Relation between genes expression was modelised using linear model. *p≤0.05; *** p≤0.001

Extended data figure 5: gluconeogenesis and glycolysis genes in human kidney biopsies classified using a computational approach.

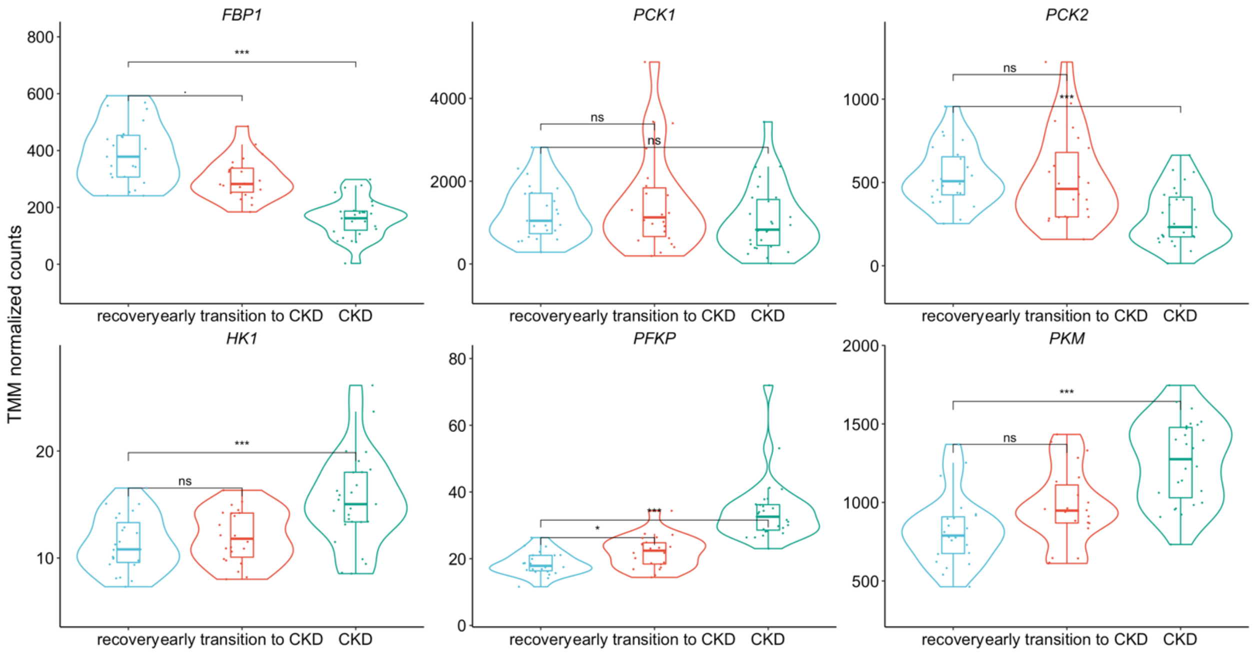


*FBP1, PCK1, PCK2, HK1, PFKP* and *PKM* genes expression data from RNAseq performed 3 and 12 months after transplantation and classified as recovery (red, n=23), early transition to CKD (green, n=22) and CKD (blue, n=27) using machine learning computational approach[12].

Extended data figure 6: FBP1 and PKM expression according to one-year eGFR

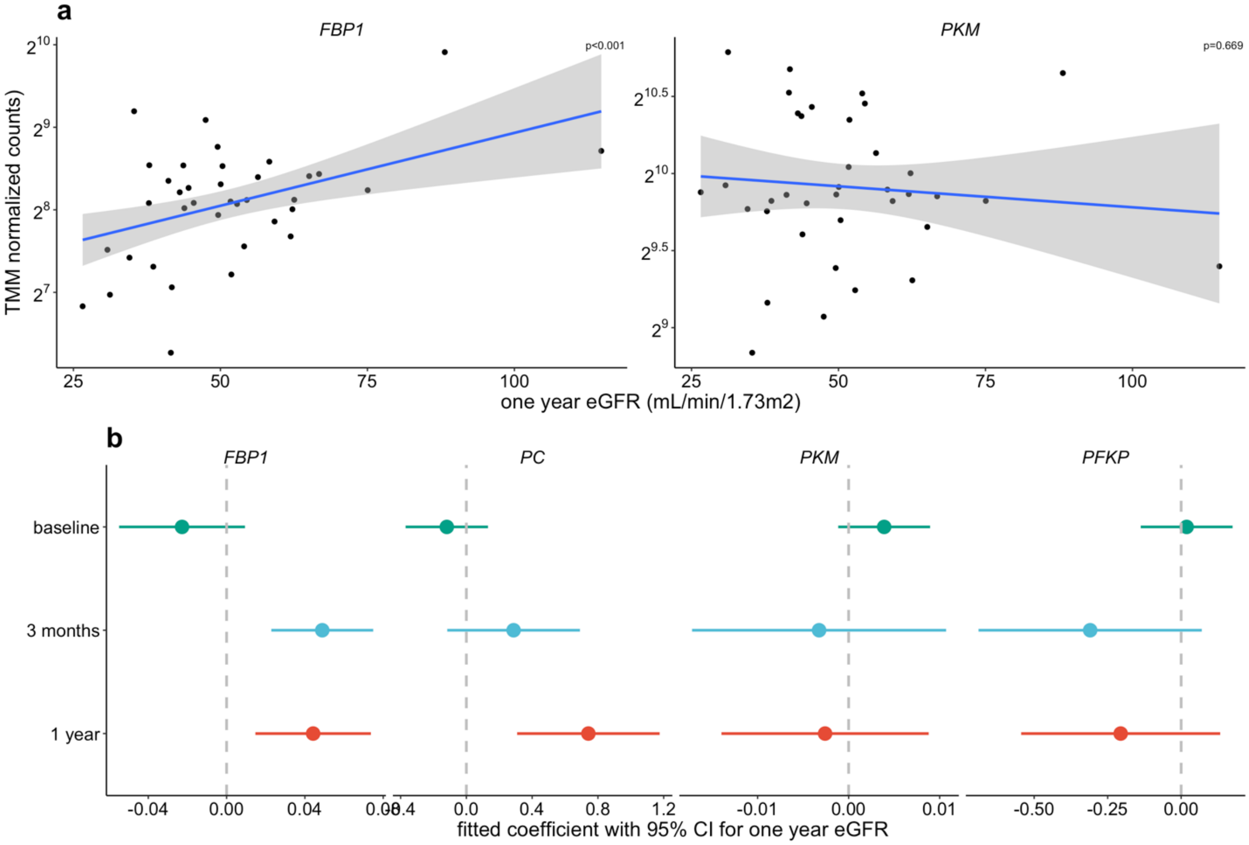


Relation between glycolytic and gluconeogenic gene expression and one-year renal function in kidney allograft recipients (a) Relation between FPB1 (left panel), *PKM* (right panel) genes expression at 3 months after transplantation and one-year GFR estimated by the CKD-EPI equation. The regression was fitted using a robust linear model. (b) Relation between one-year eGFR and *FBP1*, *PC*, *PKM* and *PFKP* gene expression at 3 different timepoints (during the transplantation, 3 and 12 months after) in kidney allograft recipients. Each dot represents the coefficient (±95% CI) of the robust linear model fitting the relation between gene expression levels and one-year eGFR. Positive value indicates a positive association.

Extended data figure 7: glucose and lactate status definition

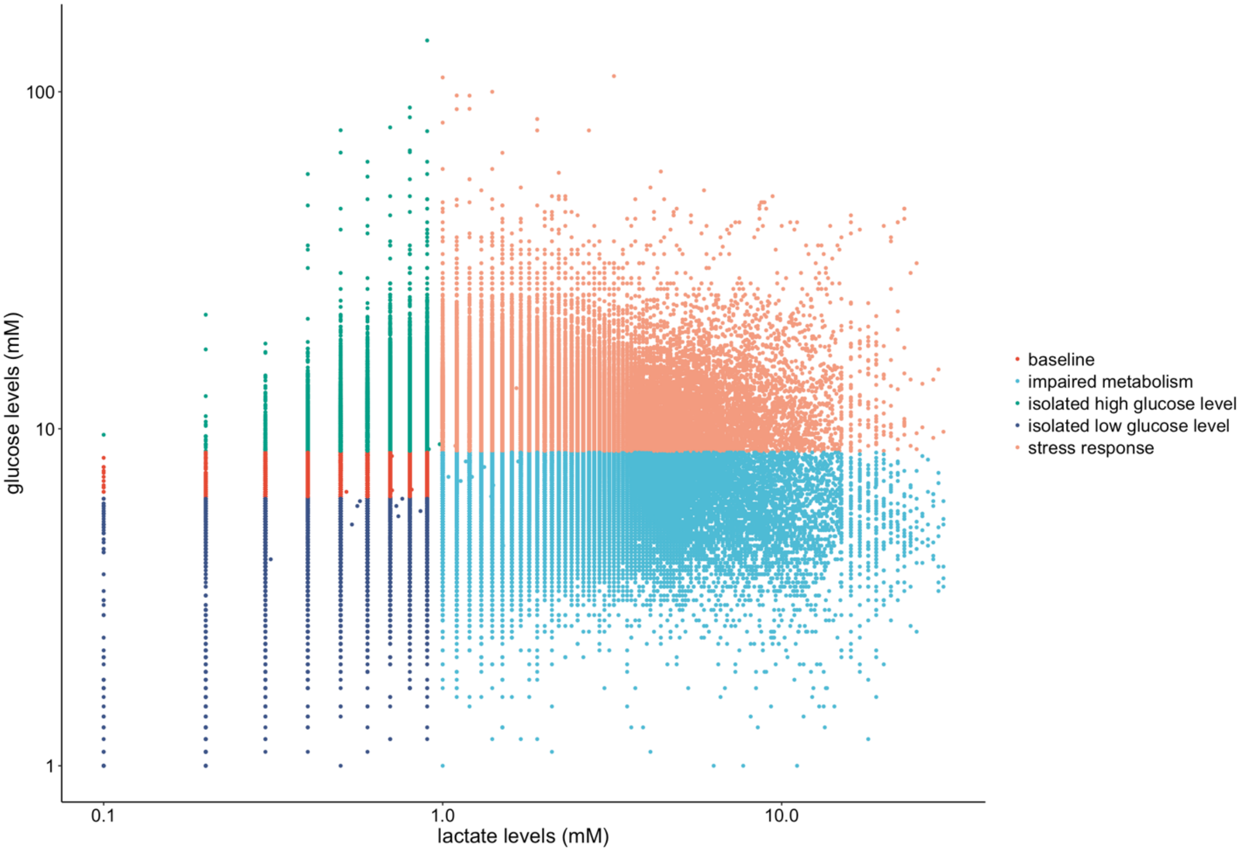


Scatter plot showing all the glucose and lactate values recorded in the ICU datasets (n=661’557). Five metabolism status were defined: baseline (lactate levels below median and with glucose levels between the 25th and the 50th percentile) ; impaired metabolism (lactate levels above the median with glucose level below the 75th percentile) ; isolated low glucose level (lactate levels below median with glucose levels above the 75th percentile) ; isolated high glucose level (lactate levels below median with glucose levels below the 25th percentile) and stress response (lactate levels above median and glucose levels above the 75th percentile).

Extended data figure 8: balance of variables for AKI before and after matching

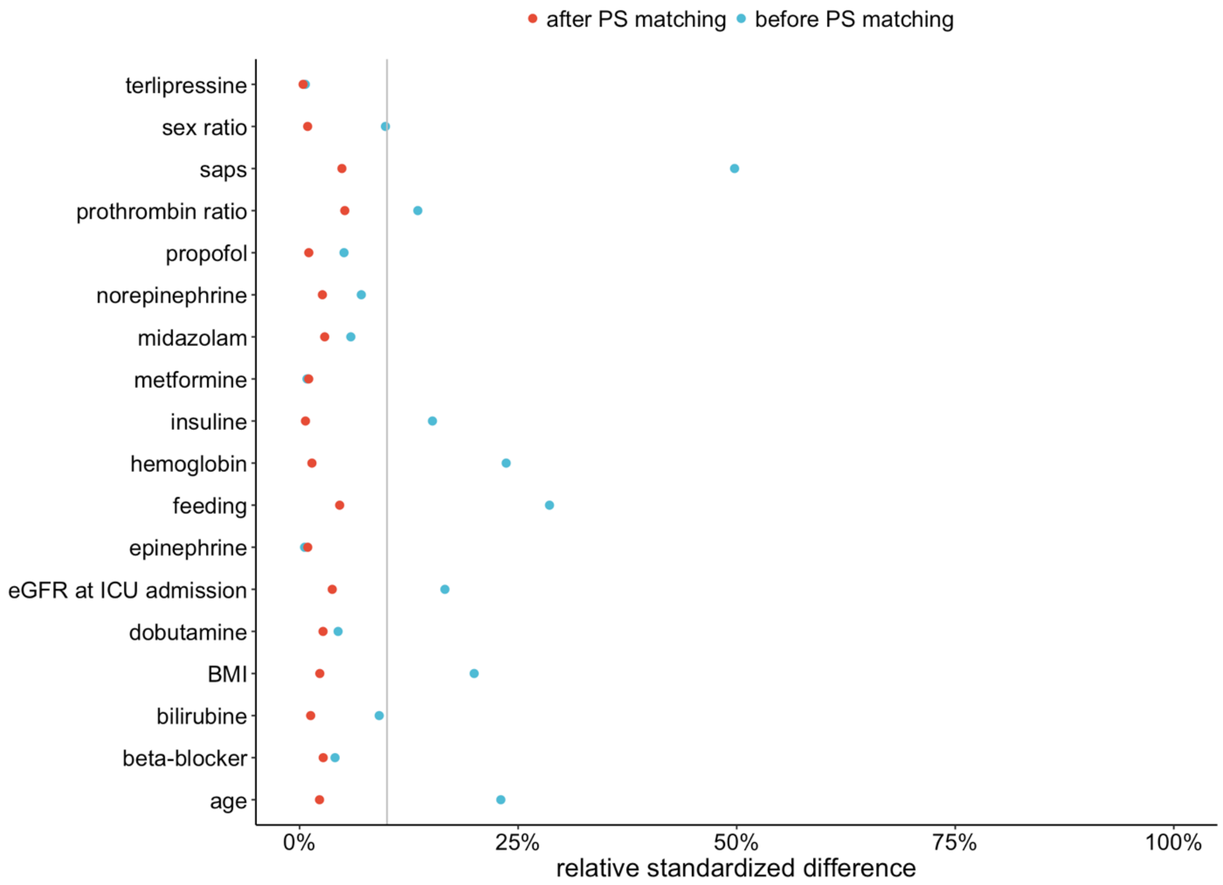


Relative standardised differences between AKI and non-AKI patients, before and after propensity score matching for each variable included in the propensity score. A relative standardised difference less than 10% was considered to support the assumption of balance between groups.

Extended data figure 9: flow chart after PSM matching after matching in the AKI dataset

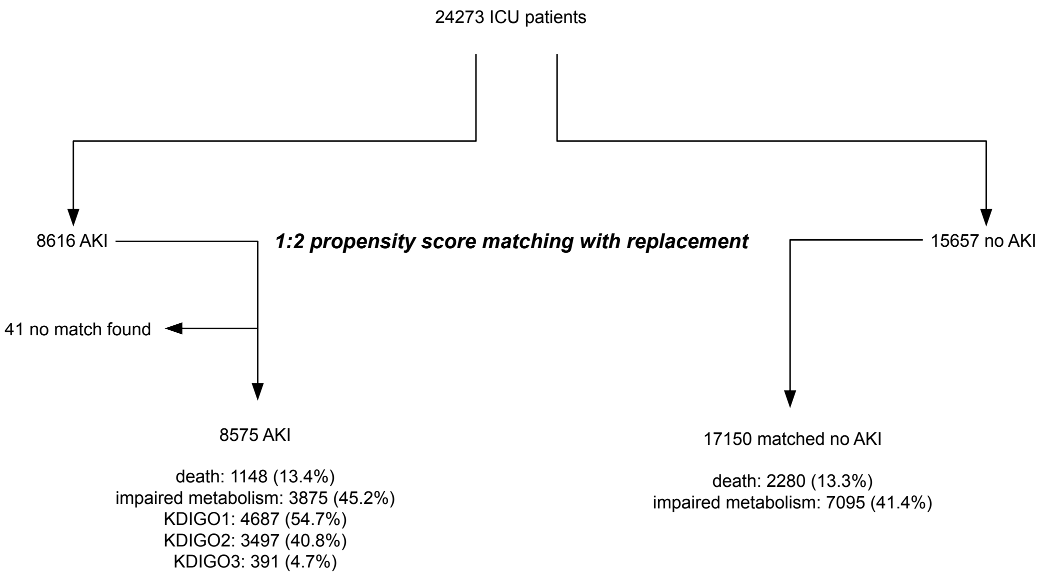


Flowchart of the propensity score matching strategy

Extended data figure 10: relation between impaired metabolism pattern rate for each kidney allograft recipient and POD15 serum creatinine.

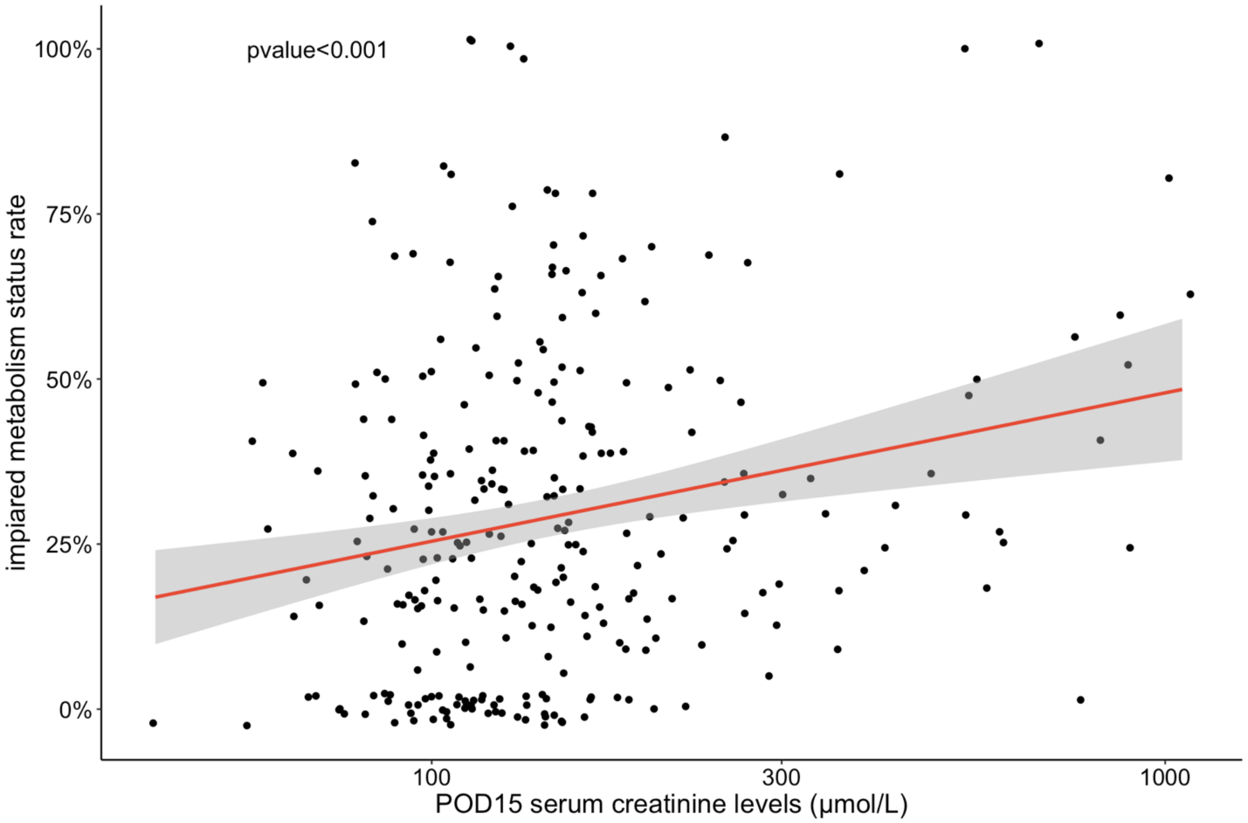


Relation between proportion of metabolism status classified as impaired metabolism, for each allograft kidney recipients during the ICU stay and the post operative day 15 serum creatinine levels, assessed by robust linear model

Extended data figure 11: ICU mortality according to KDIGO and impaired metabolism status

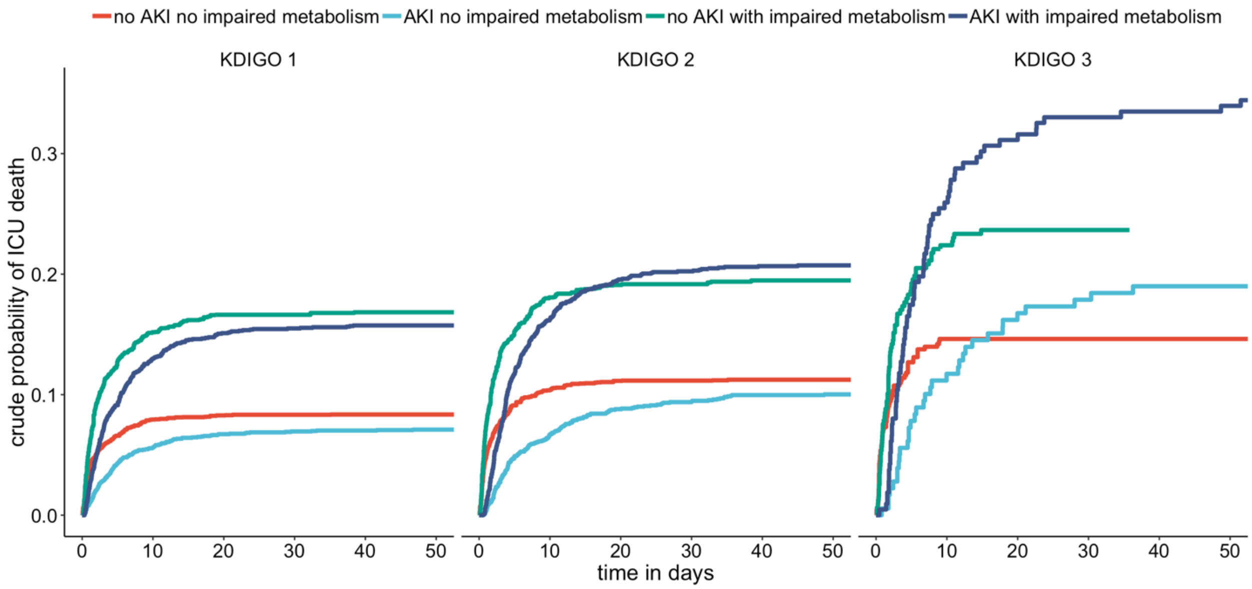


Cumulative incidence curves for ICU mortality in the cohort of ICU patients matched using a propensity score for AKI comparing 4 groups (with or without impaired metabolism pattern and with or without AKI) stratified for KDIGO.

Extended data figure 12: flow chart after matching in the thiamine dataset

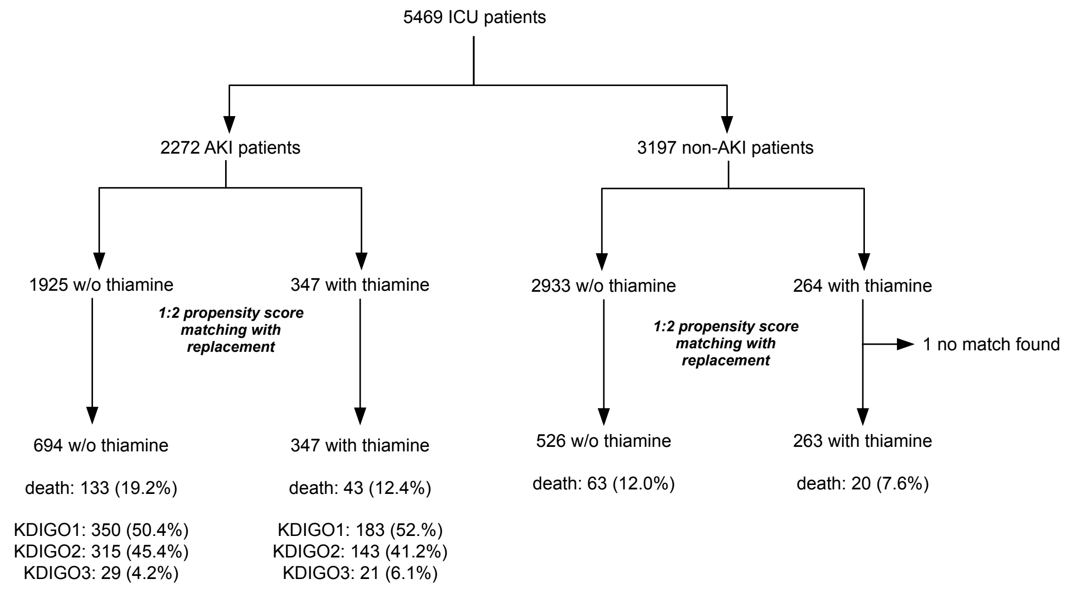


Flowchart of the propensity score matching strategy

Extended data figure 13: balance of variables before and after matching in the thiamine dataset

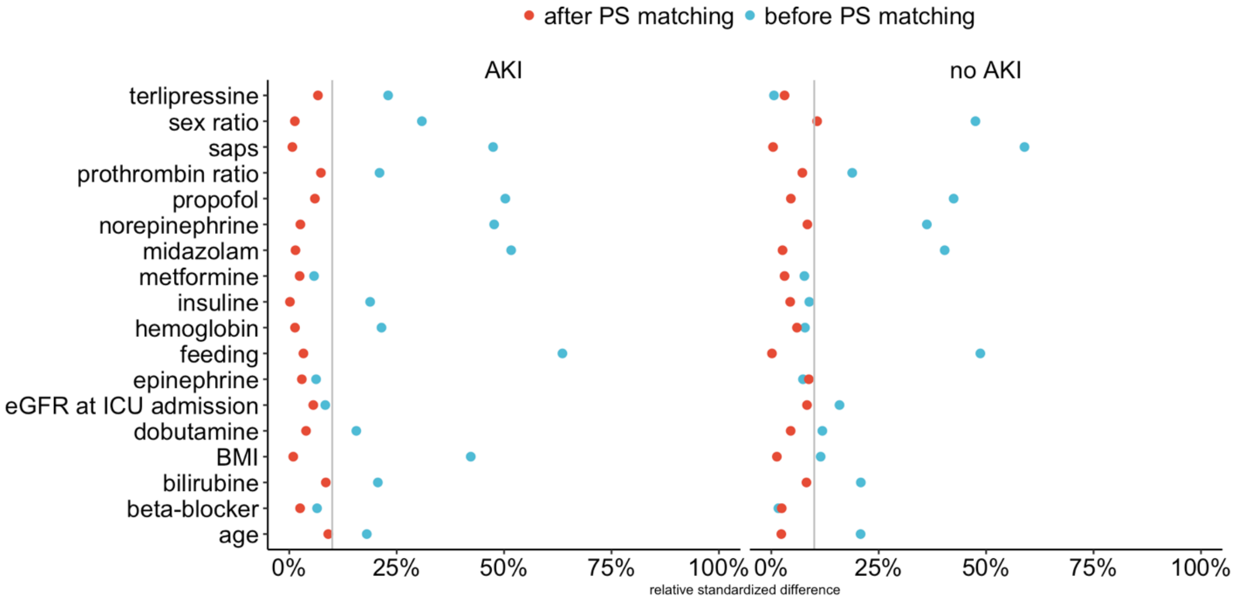


Relative standardised differences between patients receiving or not thiamine supplementation, with or without AKI, before and after propensity score matching for each variable included in the propensity score. A relative standardised difference less than 10% was considered to support the assumption of balance between groups.
